# IRE1 Regulates TOR Signaling via RIDD of *RAPTOR1b* to Coordinate Growth and Stress Adaptation

**DOI:** 10.1101/2025.10.30.685559

**Authors:** Brandon C. Reagan, Joo Yong Kim, Evan Angelos, Josh V. Vermaas, Federica Brandizzi

## Abstract

Plant growth and stress resilience depend on integrating diverse signals into coordinated cellular responses. The endoplasmic reticulum (ER) stress sensor IRE1 maintains ER homeostasis and modulates Target of Rapamycin (TOR) signaling. Here, we reveal that TOR misregulation in an *ire1ab* mutant reduces sensitivity to the stress hormone abscisic acid (ABA), mediated by TOR-dependent phosphorylation of the ABA receptor PYL1. Further, we show that IRE1’s endonuclease activity is required for TOR regulation, acting independently of the canonical IRE1/bZIP60 unfolded protein response. Instead, it occurs via Regulated IRE1- Dependent Decay (RIDD) of specific transcripts. We identify *RAPTOR1b* as a direct RIDD target, establishing a mechanistic link between ER stress sensing and TOR signaling. RIDD-mediated degradation of *RAPTOR1b* mRNA is required for appropriate ABA responses and stress adaptation. These findings uncover a noncanonical IRE1-TOR signaling axis that fine-tunes growth and stress responses through selective mRNA decay.

## Introduction

Balancing growth with adaptation to environmental stress is crucial for plant survival. Most biotic and abiotic stressors trigger proteotoxic stress in the endoplasmic reticulum (ER)^1,2^, a key hub for protein folding, lipid synthesis, and intracellular transport^3–6^. To cope with misfolded proteins, plants activate cytoprotective responses, such as the unfolded protein response (UPR)^7–9^.

A conserved component of the UPR is Inositol-Requiring Enzyme 1 (IRE1), an ER stress sensor that splices the mRNA of the transcription factor bZIP60 to activate UPR target genes^8,10–12^. In addition, IRE1 degrades specific transcripts via Regulated IRE1-Dependent Decay (RIDD)^13–16^, to modulate in autophagy, stress responses, and translational control^16–18^.

The membrane-bound transcription factor (TF), bZIP28, also contributes to ER stress tolerance^19,20^. Although single *bZIP60* or *bZIP28* mutants show no major phenotype under normal conditions, the double mutant (*bzip28/60*) exhibits WT-like growth under physiological conditions but is hypersensitive to induced ER stress,^20–22^. In contrast, the loss of two of the three IRE1 homologs (*ire1ab*) displays stunted root growth without induced stress^8,23,24^, suggesting that IRE1 regulates development through additional pathways.

IRE1 limits the activity of the Target of Rapamycin (TOR) pathway and the *ire1ab* mutant exhibits TOR hyperactivation in the root tip and impaired root elongation^24^, but how IRE1 regulates TOR remains unresolved. TOR is a conserved complex composed of TOR, RAPTOR, and LST8^25–27^. In plants, TOR influences cell division, hormone signaling, and overall growth^25,27,28^.

Abscisic acid (ABA) orchestrates abiotic stress responses through PYL receptors, and TOR inhibits ABA signaling by phosphorylating PYL1^29–33^. Although ER stress responses to heat involve IRE1-mediated bZIP60 splicing, other stressors like salt and ABA do not induce this splicing event^23,34^. Still, *IRE1* mutants are hypersensitive to salt, suggesting an alternative, bZIP60-independent, role for IRE1 in abiotic stress responses^23,35–37^.

Here, we reveal a direct mechanistic link between IRE1 and TOR signaling via RIDD-mediated regulation of *RAPTOR1b*. We show that IRE1 modulates TOR activity by degrading *RAPTOR1b* mRNA, integrating ER stress responses with ABA signaling. Our findings uncover a noncanonical IRE1–TOR axis essential for balancing growth and environmental adaptation in plants.

## Results

### The loss of IRE1 leads to decreased sensitivity to ABA

Given that IRE1 antagonizes TOR activity^24^, and TOR inhibits ABA signaling^33^, we hypothesized that loss of IRE1 impairs ABA responses. Consistent with this hypothesis, *ire1ab* exhibits a small but significant reduction in ABA content compared to wild type (Col-0, WT)^22^. To investigate whether *ire1ab* is defective in ABA responses, we examined its sensitivity to exogenous ABA at different developmental stages. First, we assessed germination on ABA- containing media^38,39^. While only ∼30% of Col-0 seeds germinated after 3 days in ABA, *ire1ab* seeds showed significantly higher germination (∼60%) (Figure 1A, B), indicating reduced ABA sensitivity. Next, we evaluated root growth responses to ABA. Col-0 and *ire1ab* seeds were germinated on ½ LS media, then transferred at day 3 to media with 1μM or 10μM ABA for 7 days. As expected^40,41^, ABA reduced primary root growth in Col-0, but had significantly lower effect in *ire1ab*, and at 1μM ABA, *ire1ab* grew longer than mock control (Figure 1C, D). Finally, expression of ABA-responsive genes (*RD29A*, *RD29B*, *RAB18*) in *ire1ab* root tips was significantly lower than in Col-0 after 6 h of ABA treatment (Figure 1E). These findings demonstrate that *ire1ab* is less responsive to exogenous ABA, revealing a novel role of IRE1 in ABA signaling.

**Figure 1:**
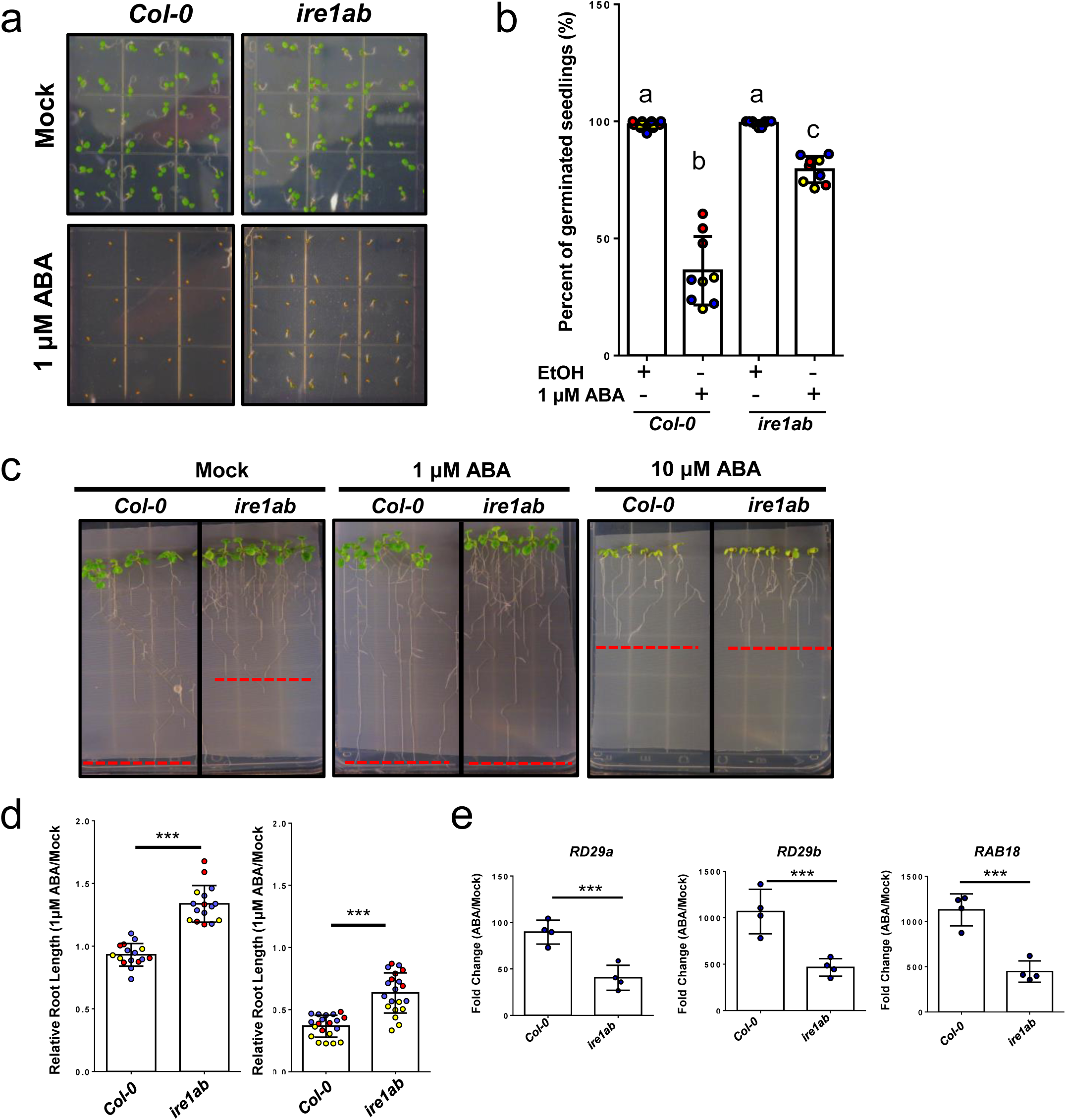
*Ire1ab* exhibits decreased sensitivity to ABA. **a.** Representative images of *Col-0* and *ire1ab* seeds germinated on plates with 1μM ABA or mock treatment. **b.** Quantification of germination rates at 3 days across three biological replicates (replicate indicated by circle color). Each data point represents the germination rate of 28 seeds on one plate for each biological replicate. **c.** Representative images of 10-day-old seedlings grown on plates with 0 (Mock), 1μM, or 10μM ABA. Red dashed lines depict the average root length for each genotype under each treatment. **d.** Quantification of relative root lengths (ABA/Mock) for 10-day-old seedlings (7 days post transfer) on 1μM (left) and 10μM (right) ABA treatment across three biological replicates (replicate indicated by circle color). Each data point represents the average root length of 10 seedlings on one plate (one technical replicate). **e.** qRT-PCR analysis of ABA- responsive gene expression in Col-0 and *ire1ab* seedlings. Graphs depicting the change in expression (ABA/mock) across four biological replicates. Each data point represents the average of 3 technical replicates. **b,d, and e.** Asterisks indicate statistically significant differences by unpaired 2-tailed Student’s t-test. **b**. Letters indicate groups by statistical significance using two-way ANOVA.

### Inhibition of TOR restores ABA sensitivity in ire1ab

The short root phenotype of *ire1ab* is caused by the hyperactivation of TOR and is reversed by the TOR-specific inhibitor AZD-8055 (AZD)^24^. To investigate whether TOR misregulation in *ire1ab* contributed to the decreased sensitivity to ABA, seeds were sown on AZD, ABA, AZD and ABA, or mock plates. AZD treatment alone did not affect seed germination rates after 3 days (Figure 2A, B). Col-0 seedlings treated with both ABA and AZD germinated at similar rates compared to ABA alone (Figure 2A, B). In contrast, *ire1ab* seeds treated with both ABA and AZD exhibited a significant reduction in germination rate compared to ABA-only treatment, with a germination rate comparable to Col-0 (Figure 2A, B). These results indicate that the decreased sensitivity of *ire1ab* to ABA is due to TOR misregulation. To determine whether TOR inhibition restored ABA sensitivity in *ire1ab* during primary root growth, Col-0 and *ire1ab* seedlings were transferred to plates with AZD, ABA, AZD and ABA, or mock treatment. After 7 days of growth, low concentrations of either AZD or ABA rescued the short-root phenotype of *ire1ab,* while the combined treatment restored ABA sensitivity in *ire1ab* (Figure 2C, D).

**Figure 2:**
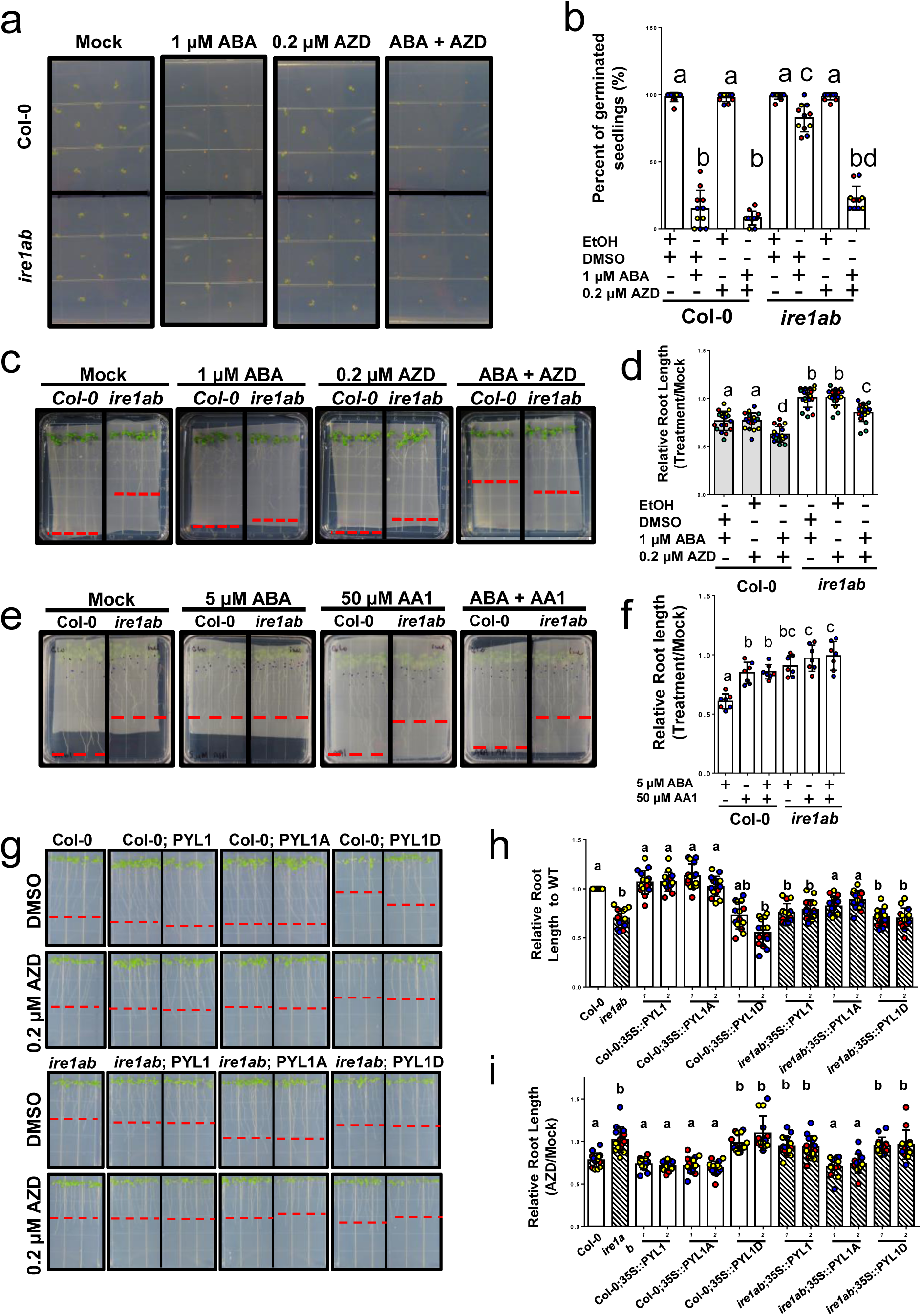
Hyperactivation of TOR leads to inactivation of ABA receptors in *ire1ab*. **a.** Representative images of *Col-0* and *ire1ab* seeds germinated on plates with 1 μM ABA, 0.2 μM AZD, ABA and AZD, or mock treatment. **b.** Quantification of germination rates at 3 days old across three biological replicates (replicate indicated by circle color). Each data point represents the germination rate of 28 seeds on one plate for each technical replicate. **c.** Representative images of 10-day-old seedlings grown on plates with 1 μM ABA, 0.2 μM AZD, ABA and AZD, or mock treatment. Red dashed lines depict the average root length for each genotype under each treatment. **d.** Quantification of relative root lengths (Treatment/Mock) for 10-day-old seedlings (7 days post transfer) on 1 μM ABA, 0.2 μM AZD, ABA and AZD, or mock treatment plates across three biological replicates (replicate indicated by circle color). Each data point represents the average root length of 10 seedlings on one plate for each technical replicate **e.** Representative images of 10 day old seedlings (7 days post transfer) on plates with 5μM ABA, 50μM AA1, ABA and AA1, or mock treatment. The red dashed line represents the average root length for each genotype. **f.** Quantification of relative root length for 10-day old seedlings on 5μM ABA, 50μM AA1, ABA and AA1, or mock treatment across three biological replicates (replicate indicated by circle color). Each data point represents the average root length of 10 seedlings on one plate (one technical replicate). **g.** Representative images of Col-0 and *ire1ab* seedlings along with two independent lines of 35S::PYL1, 35S::PYL1A, and 35S::PYL1D in both Col-0 and *ire1ab* backgrounds grown on ½ LS plates without sucrose and supplemented with 0.2μM AZD or mock treatment. The red dashed line represents the average root length for each genotype. **h-i.** Quantification of relative root length (normalized to Col-0 root length) (h) and (AZD/Mock treatment) **(i)** for 10 day old seedlings across three biological replicates (replicate indicated by circle color). Each data point represents the average root length of 10 seedlings on one plate (one technical replicate). The bar pattern represents the genetic background for each line (Col- 0, plain) or (*ire1ab,* slashed) **b, d, f, h and i.** Statistical significance was determined by two-way ANOVA with multiple comparisons. Letters indicate groups by statistical significance.

These findings demonstrate that TOR hyperactivation in *ire1ab* underlies the reduced sensitivity to ABA across multiple stages of plant growth and development.

### Hyperactivation of TOR in ire1ab inactivates PYL1

To investigate if ABA signaling is disrupted at the receptor level in *ire1ab*, seedlings were treated with the ABA receptor inhibitor, AA1^42^. Col-0 seedlings grown in the presence of AA1 exhibited a reduction in primary root length, as previously reported^42^, while *ire1ab* was unaffected (Figure 2E, F). Combined treatment with ABA and AA1 blocked ABA-dependent inhibition of primary root growth in Col-0 but had no effect on *ire1ab*, suggesting that ABA receptors in *ire1ab* may be inactive.

To determine if PYL1 was differentially phosphorylated at the TOR-specific phosphorylation site (S119)^33^ in *ire1ab* seedlings, root tips were harvested from Col-0 and *ire1ab* seedlings for targeted mass spectrometry. A synthetic peptide served as a standard for detecting phosphorylated PYL1. A representative spectrum confirming full coverage of the targeted site is shown in Supplemental Figure 1A. In two of the three *ire1ab* samples, PYL1S119 was highly phosphorylated (Supplemental Figure 1B), whereas no phosphorylated peptide was detected in Col-0 samples. This indicates that PYL1 is active in Col-0 root tips but inactivated in *ire1ab*. Sequence constraints prevented testing for phosphorylation of other PYLs, but this site for TOR phosphorylation is conserved across all 14 PYL receptors (Supplemental Figure 1C).

To assess the impact of constitutive PYL1 phosphorylation on plant growth, transgenic lines were generated to express WT (PYL1S119), phospho-null (PYL1S119A), and phospho-mimic (PYL1S119D) versions of PYL1 under the control of the CaMV35S promoter. Two independent lines of each construct in WT and *ire1ab* backgrounds were selected for analysis. Expression of either WT PYL1 or PYL1S119A in the Col-0 did not affect primary root length, while expression of PYL1S119D in induced a short-root phenotype comparable to that of the *ire1ab* mutant (Figure 2G, H). Moreover, PYL1S119D expression reduced sensitivity to TOR inhibition by AZD (Figure 2G, I). In contrast, the expression of PYL1S119A in *ire1ab* partially rescued the short-root phenotype and restored sensitivity to AZD treatment while expression of PYL1S119D had minimal impact on root length and did not alter AZD sensitivity compared to untransformed *ire1ab* seedlings (Figure 2G-I).

These results are consistent with the hypothesis that the TOR-mediated inhibition of ABA receptors contributes to the short-root phenotype of *ire1ab*.

### Hyperactivation of TOR in ire1ab is independent of the canonical UPR

To gain a deeper understanding of the connection between IRE1 and TOR, we investigated other core UPR mutants for changes in TOR activity and ABA responses. When treated with AZD, *bzip28/60* seedlings exhibited a reduction in primary root length comparable to Col-0 seedlings (Figure 3A, B). Additionally, *bzip28/60* root tips exhibited WT levels of S6K phosphorylation indicating that TOR is not hyperactivated (Figure 3C, D). Furthermore, we treated *bzip28/60* with exogenous ABA to evaluate whether ABA signaling was compromised and found that exogenous ABA inhibited the primary root growth of *bzip28*/*60* seedlings similar to Col-0 (Figure 3E, F). Collectively, these findings indicate that the link between IRE1 and TOR is independent of the canonical UPR-TFs.

**Figure 3:**
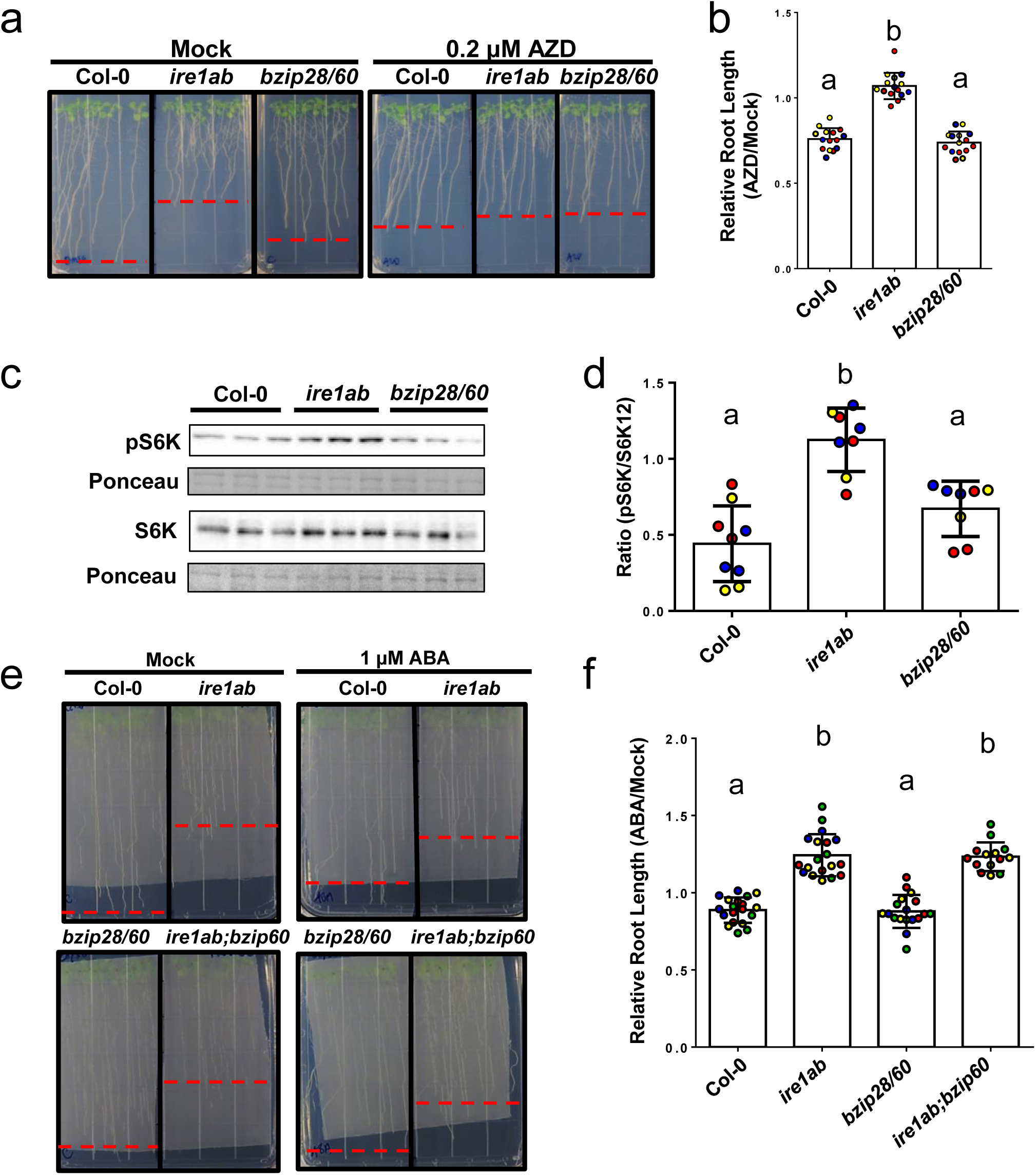
IRE1-dependent TOR regulation is independent of canonical ER stress pathways. **a.** Representative images of 10-day-old Col-0, *ire1ab*, and *bzip28/60* seedlings grown on plates supplemented with 0.2μM AZD or mock treatment. The red dashed line represents the average root length for each genotype. **b.** Quantification of relative root length (AZD/Mock) for AZD-treated seedlings across three biological replicates (circle color represents each independent biological replicate). Each data point represents the average root length of 10 seedlings on one plate for each technical replicate. **c.** Representative western blots of S6K phosphorylation levels in Col-0, *ire1ab,* and *bzip28/60* root tips using primary antibodies raised against S6K1/2 (bottom blot) and phosphorylated form of S6K (Top blot). Ponceau staining of the membrane was used as loading control. **d.** Quantification of the ratio of S6K and pS6K band intensity for three independent biological replicates, indicated by circles. **e.** Representative images of 10-day-old Col-0, *ire1ab, bzip28/60,* and *ire1ab;bzip60* grown on mesh on ½ LS plates supplemented with 1% sucrose and either 1 μM ABA or mock conditions. The red dashed line represents the average root length for each genotype. **f.** Quantification of relative root length (ABA/Mock) for ABA treated seedlings across three biological replicates (circle color represents each independent biological replicate). Each data point represents the average root length of 10 seedlings on one plate for each technical replicate. **b and d.** Statistical significance was determined by Student’s t-test. **f.** Statistical significance was determined by 2-way ANOVA. Letters indicate groups by statistical significance.

### The endonuclease activity of ire1ab is involved in IRE1-dependent TOR regulation

While IRE1’s endonuclease activity is primarily known for the splicing of *bZIP60* mRNA, it also regulates transcript abundance via RIDD^14,17,18^. *In vitro* RNase activity assays demonstrated that 4μ8C, a specific inhibitor of IRE1 endonuclease activity^43,44^, effectively blocks the activity of Arabidopsis IRE1 (Supplemental Figure 2A-C). When Col-0 seedlings were grown on media with 4μ8C, primary root growth was inhibited, and they developed a short root phenotype similar to untreated *ire1ab* (Figure 4A, B, Supplemental Figure 2D,E). They also exhibited elevated levels of phosphorylated S6K, indicating increased TOR activity (Figure 4C, D) and supporting that IRE1 regulates TOR via its endonuclease activity. As expected, *bzip28/60* seedlings were hypersensitive to inhibition of IRE1 endonuclease activity (Figure 4A, B). To determine if the hypersensitivity of Col-0 and *bzip28/60* was due to 4μ8C-induced TOR hyperactivation, seeds were grown on media supplemented with 4μ8C and/or AZD (Figure 4E, F). AZD treatment decreased 4μ8C sensitivity in both backgrounds (Figure 4E, F).

**Figure 4:**
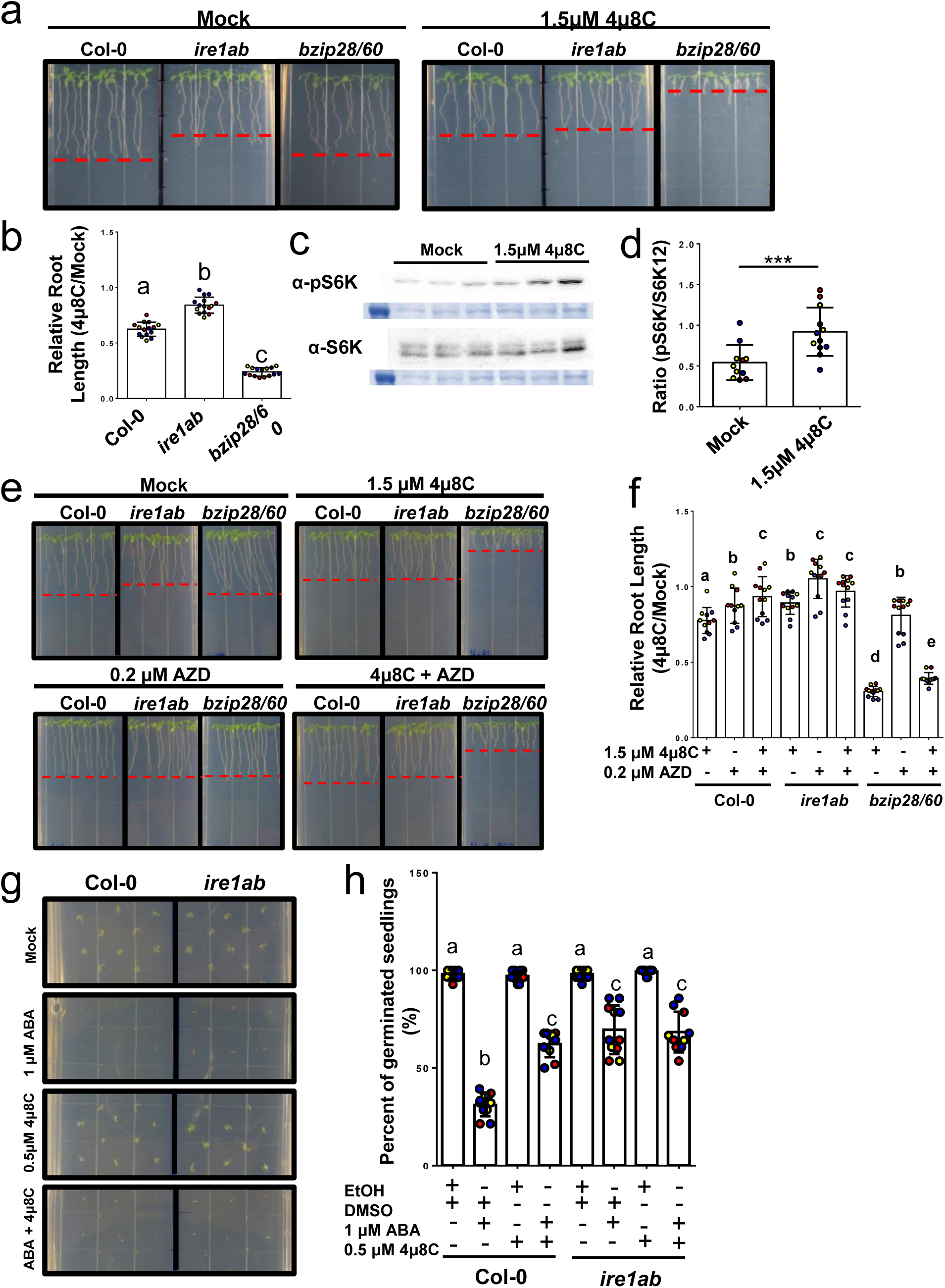
IRE1 regulates TOR via RIDD. **a.** Representative images of 10-day-old Col-0, *ire1ab,* and *bzip28/60* seedlings grown on plates supplemented with 1.5 μM 4μ8C or DMSO (mock). The red dashed line represents the average root length for each genotype. **b.** Quantification of relative root length (4μ8C/Mock) for 10-day-old seedlings across three biological replicates (indicated by circle color). Each data point represents the average root length of 10 seedlings from one plate (one technical replicate). **c.** Representative western blots showing S6K phosphorylation levels of Col-0 seedlings grown on plates supplemented with 1.5 μM 4μ8C. **d.** Quantification of the ratio of S6K and pS6K band intensity across three biological replicates. **e.** Representative images of 10-day-old Col-0, *ire1ab,* and *bzip28/60* seedlings grown on plates supplemented with 1.5 μM 4μ8C, 0.2 μM AZD, 4μ8C and AZD, or DMSO for mock treatment. The red dashed line represents the average root length for each genotype. **f.** Quantification of relative root length (Treatment/Mock) for 10-day-old seedlings across three biological replicates (indicated by circle color). Each data point represents the average root length of 10 seedlings from one plate (one technical replicate). g. Representative images of Col-0 and *ire1ab* seeds germinated on plates supplemented with 1 μM ABA, 0.5 μM 4μ8C, ABA and 4μ8C, or EtOH and DMSO mock treatment. The red dashed line represents the average root length for each genotype. **h.** Quantification of germination rates at 3 days after germination across three biological replicates (replicate indicated by circle color). Each data point represents the germination rate of 28 seeds on one plate for each biological replicate. Statistical significance was determined by a one-way ANOVA (**b**), two-tailed Student’s t-test (**d**), and two-way ANOVA (**f and h**). Letters indicate groups by statistical significance.

These results indicate that RIDD is critical for physiological plant growth and development, extending its role beyond ER stress.

Next, to assess whether disruption of IRE1 endonuclease activity affected ABA sensitivity, Col-0 and *ire1ab* seeds were germinated on ABA, 4μ8C, ABA and 4μ8C, or a mock control media. 4μ8C treatment alone did not affect germination rates for either Col-0 or *ire1ab* seeds (Figure 4G, H). However, the combined treatment resulted in reduced ABA sensitivity in Col-0, germinating at rates comparable to *ire1ab* (Figure 4G, H).

These findings further suggest that RIDD plays a significant role in regulating ABA responses through TOR.

### RAPTOR1b is a target of RIDD

Several RIDD targets involved in ER stress responses have been identified^17,18^; however, none have been linked to physiological growth. Metazoan IRE1’s RIDD targets mRNAs contain the consensus sequence (CNGCAGN) in hairpin structures^45,46^. gRIDD is a program designed to identify RIDD targets based on the presence of the mammalian consensus sequence^47^. To explore potential RIDD targets in Arabidopsis, we analyzed the transcriptome using gRIDD and identified 1,170 candidates matching the consensus sequence, along with 1,889 and 2,214 candidates for two alternative sequence variants (Supplemental data 1). Comparison of these predictions with known RIDD targets revealed that PR-4 (*AT3G04720*)^18^ and GLH19 (*AT2G43610*)^17^ contain an exact match, while PR-14 (*AT5G01870*)^17^ contained a variant (var5) sequence. The analysis also identified *RAPTOR1a* and *RAPTOR1b* as potential targets of RIDD. *RAPTOR1b* mRNA contained an exact match for the consensus sequence located within a predicted stem-loop structure, while *RAPTOR1a* mRNA^48^ contained the var5 form.

Next, we used AlphaFold3^49^ to model the interaction between IRE1B and a portion of *RAPTOR1b* mRNA (Supplemental Figure 3A). We modeled segments of *RAPTOR1b* mRNA with the WT (GCTGCAGAC) and a mutated (GCGGCGGAT) version of the consensus sequence, and focused on the KEN domain, which is crucial for IRE1’s endoribonuclease activity^50^. Modeling both WT and mutant sequences with the dimeric IRE1B structure yielded a range of potential complexes (Supplemental Figure 3A, Supplemental Data 2). Asparagine 826 (N826), located in the KEN domain, is likely involved in RNase activity^50^; so we expected that a functional complex would position N826 near the target sequence. We found that in 8 of 15 predicted complexes with the native sequence, the heavy atoms of the mRNA were within 5 Å of N826, compared with only 2 of 15 for the mutant. While AlphaFold probability predictions do not fully reflect the underlying energy landscape^51^, and RNA modeling remains challenging, the results support that mutations in *RAPTOR1b* mRNA disrupt association with the IRE1 active site. Next, we analyzed mRNA structural changes by examining θ and η angles along the backbone, analogous to Ramachandran angles in proteins^52,53^. AlphaFold3 predicted largely equivalent structures, with differences localized near altered nucleobases (Supplemental Figure 3B). Visualizations showed different bases exposed rather than base-paired. We quantified these differences using solvent-accessible surface area (SASA), finding localized changes near the mutation site (gray region in Supplemental Figure 3C). In the WT sequence, nucleobases 109-116 were more exposed, while upstream nucleobases were more exposed in the mutant sequence. Representative models (Supplemental Figure 3D) illustrate these differences, supporting the hypothesis that *RAPTOR1b* mRNA contains a putative RIDD consensus sequence recognized by the IRE1B KEN domain.

To experimentally validate *RAPTOR1b* as a RIDD target, we treated seedlings with Actinomycin D (ActD) to block transcription and harvested root tips to monitor *RAPTOR1b* mRNA degradation via qRT-PCR. In Col-0, *RAPTOR1b* mRNA abundance decreased by 50-60% after 12 hours of treatment (Figure 5A). In contrast, degradation was significantly reduced (∼20%) in *ire1ab* and was comparable to that of the transcript of the TORC component, *LST8-1*, which lacks a RIDD consensus sequence. Additionally, *RAPTOR1b* mRNA degradation by IRE1B was measured using *in vitro* RNase assays with the purified cytosolic domain of IRE1B (IRE1B_­­_). Total RNA was incubated with and without IRE1B_­­_or a heat-inactivated IRE1B_­­_(IRE1B_­­_). qRT-PCR was used to measure transcript abundance of unspliced *bZIP60* (*ubZIP60*; positive control), *RAPTOR1b*, and *LST8-1* (negative control). As expected following incubation with IRE1B_­­_, *ubZIP60* transcript abundance was largely reduced (>95%), while incubation with IRE1B_­­_only led to a slight reduction in abundance (Figure 5B). Similalry, *RAPTOR1b* transcript abundance also exhibited a strong reduction (80%) following incubation with IRE1B_­­_but not with IRE1B_­­_. In contrast, LST8-1 transcript abundance remained unchanged following incubation with IRE1B_­­_, indicating that the reduction in *RAPTOR1b* mRNA is specific and IRE1B-dependent (Figure 5B).

**Figure 5:**
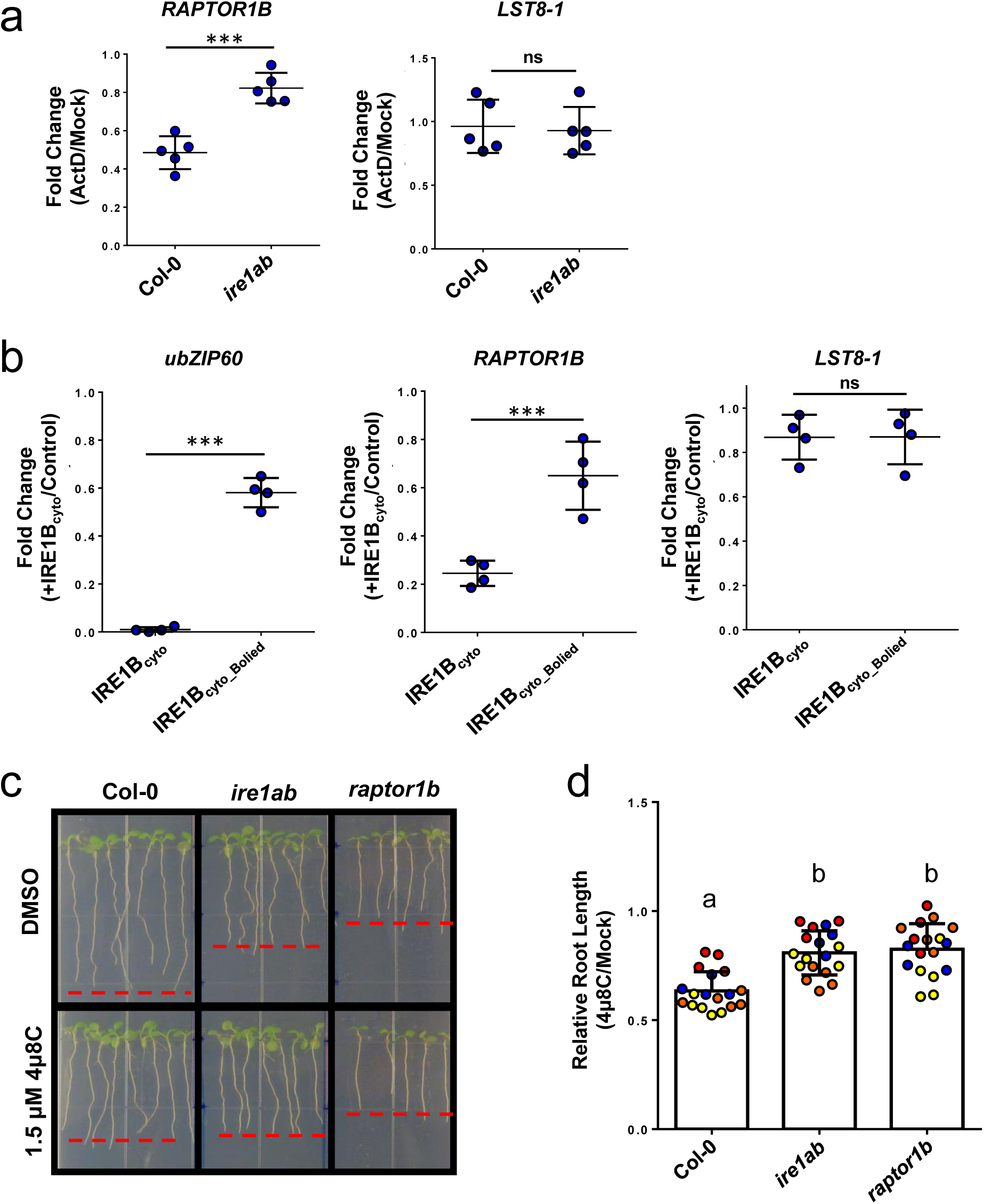
RAPTOR1b is a target of RIDD under normal growth conditions. **a.** qRT-PCR analysis of RAPTOR1b and LST8-1 (negative control) transcript abundance in the root tips of 10-day-old Col-0 and ire1ab seedlings following 12 hours of treatment with 25μM ActD across five biological replicates. Each data point represents the average of three technical replicates **b.** qRT-PCR analysis of ubZIP60, RAPTOR1b, and LST8-1 transcript abundance following in vitro RNA degradation assays with IRE1bcyto or heat-inactivated IRE1bcyto_boiled. Each data point represents the average of three technical replicates across four biological replicates. **c.** Representative images of 10-day-old Col-0, ire1ab, and raptor1b seedlings grown on plates supplemented with 1.5μM 4μ8C or DMSO as a mock treatment. Red dashed lines represent the average root length of each genotype under each treatment. **d**. Quantification of relative root length (4μ8C/Mock) across three biological replicates (indicated by circle color). Each data point represents the average root length of 10 seedlings from one plate (one technical replicate). Statistical significance was determined by two-tailed student t-tests (**a-b**) and one way ANOVA (**d**). Letters denote groups by statistical significance.

We next aimed to test whether IRE1-dependent TOR hyperactivation was mediated by misregulation of *RAPTOR1b* abundance using an established *raptor1b* KO line (*SALK_078159*)^48^. *Raptor1b* seedlings exhibited short primary root and reduced growth compared to Col-0 as well as reduced sensitivity to IRE1 nuclease inhibition, suggesting that *RAPTOR1b* mRNA degradation is critical for IRE1-dependend TOR regulation (Figure 5C, D).

Together, these results support that *RAPTOR1b* mRNA is a RIDD target in physiological conditions of growth, and that impaired *RAPTOR1b* mRNA degradation contributes to TOR hyperactivation in *ire1ab*. Notably, the *RAPTOR1b* consensus sequence is conserved across plant species (Supplemental Figure 4), suggesting that RIDD-mediated regulation of TOR may be a common feature in plants.

### RIDD of RAPTOR1b is required for proper responses to salt stress

Because previous studies linked hypersensitivity to salt stress and TOR hyperactivation to defects in meristematic cell development in *ire1ab*^24,37^, we hypothesized that salt hypersensitivity may be related to TOR misregulation in *ire1ab*. As previously reported^37^, we found that *ire1ab* was hypersensitive to NaCl (Supplemental Figure 5). *bzip28/60* did not exhibit hypersensitivity to salt stress, suggesting that the phenotype is specific to *ire1ab* and is not due to a general UPR deficiency (Figure 6A, B). We next tested whether misregulation of TOR activation contributed to the salt hypersensitivity in *ire1ab*. Seeds were sown on NaCl, AZD, NaCl and AZD, or mock plates and TOR inhibition completely rescued the short-root phenotype of *ire1ab* on salt plates, indicating that TOR misregulation drives the hypersensitivity of *ire1ab* to salt stress (Figure 6C, D).

**Figure 6:**
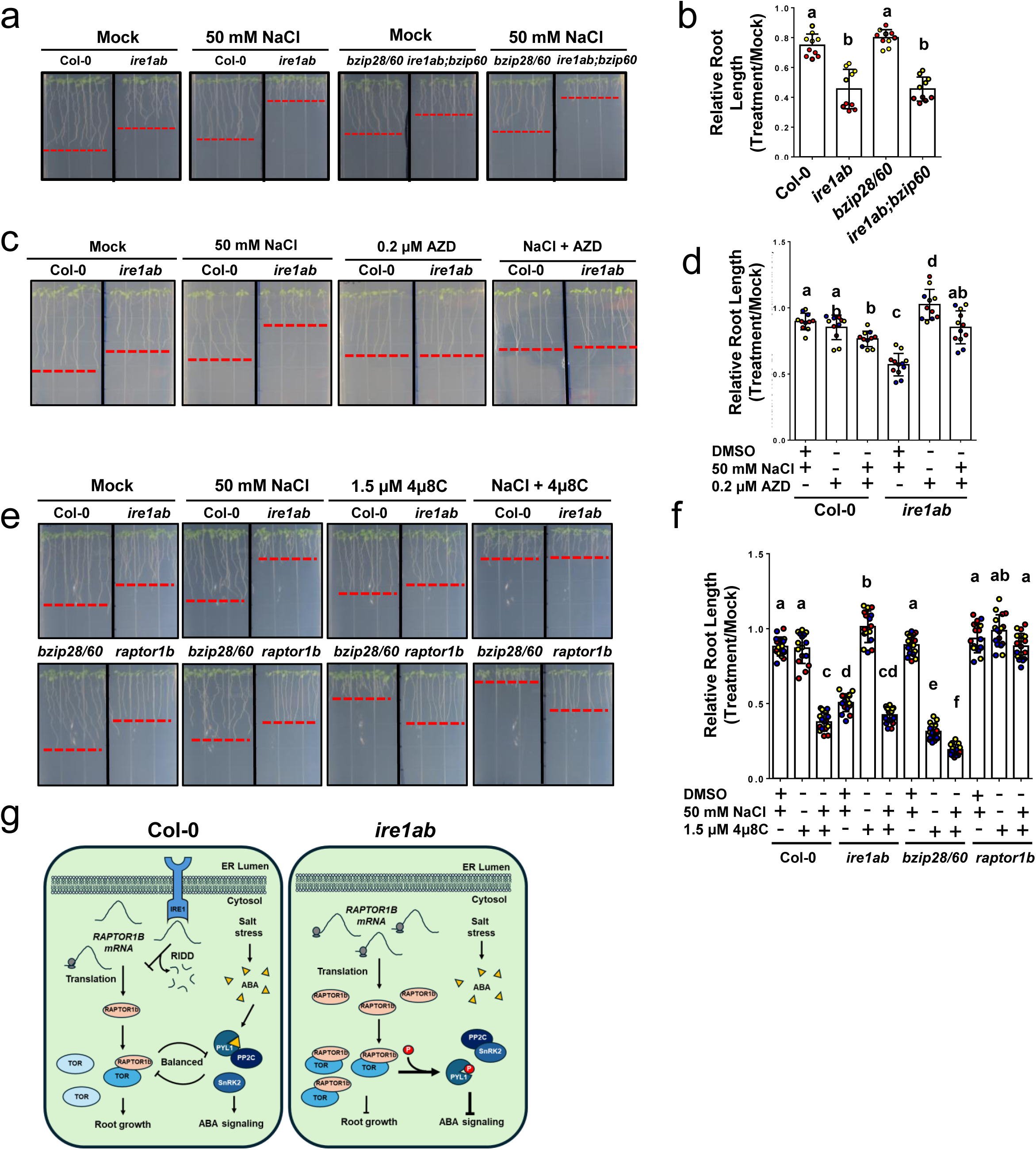
RAPTOR1b transcript degradation via RIDD is required for ABA response to salt stress. **a.** Representative images of Col-0, *ire1ab, bzip28/60,* and *ire1ab;bzip60* seedlings grown on plates with or without 50 mM NaCl. Red dashed lines represent the average root length of each genotype on each treatment. **b.** Quantification of relative root length (NaCl/Mock) across three biological replicates (indicated by circle color). Each data point represents the average root length of 10 seedlings from one plate (one technical replicate). **C.** Representative images of Col-0 and *ire1ab* seedlings on ½ LS plates supplemented with 50 mM NaCl, 0.2μM AZD, NaCl and AZD, or DMSO for mock control treatment. Red dashed lines represent the average root length of each genotype under each treatment. **d.** Quantification of relative root length (Treatment/Mock) across three biological replicates (indicated by circle color). Each data point represents the average root length of 10 seedlings from one plate (one technical replicate). **e.** Representative images of Col-0, *ire1ab, bzip28/60,* and *raptor1b* seedlings grown on plates supplemented with 50 mM NaCl, 1.5 μM 4μ8C, NaCl and 4μ8C, or mock control. Red dashed lines represent the average root length of each genotype for each treatment. **f.** Quantification of relative root length (treatment/mock) across three biological replicates (indicated by circle color). Each data point represents the average root length of 10 seedlings from one plate in one technical replicate. **g.** Model of IRE1-dependent regulation of TOR and ABA signaling responses via RIDD of *RAPTOR1b* mRNA. During WT plant growth and development in physiological conditions (Left), IRE1 regulates the translation of *RAPTOR1b* mRNA via RIDD to establish a balanced pool of RAPTOR1b protein and maintain TOR activity. The TOR complex forms a reciprocal regulatory loop with ABA signaling machinery, establishing a balance between growth and ABA responses. Under abiotic stress conditions, such as salt stress, the balance can shift towards TOR inhibition, allowing for increased ABA signaling. In the *ire1ab* mutant (Right), *RAPTOR1b* transcript abundance is elevated due to loss of RIDD. This leads to increased RAPTOR1b protein abundance and increased activation of TOR in the root tip. This increased TOR activation results in inhibition of ABA signaling through the phosphorylation of PYL1. This inhibition shifts the balance away from ABA stress responses and towards unchecked growth, leading to compromised responses to ABA-dependent stresses such as salt stress. Statistical significance was determined by one-way ANOVA **(b)** or two-way ANOVA **(d and f). b, d,** and **f,** Letters indicate groups by statistical significance.

To determine if RIDD-mediated degradation of *RAPTOR1b* mRNA is required for proper salt stress responses, we grew Col-0, *ire1ab, bzip28/60,* and *raptor1b* seedlings on NaCl, 4μ8C, NaCl and 4μ8C, or mock plates. Col-0 and *bzip28/60* seedlings exhibited similar reductions in primary root length under salt stress alone while the combined treatment led to hypersensitivity similar to *ire1ab* (Figure 6E, F). In contrast, *raptor1b* seedlings did not exhibit a difference in root length between salt plates and the combined treatment, indicating that the salt stress hypersensitivity in *ire1ab* depends on *RAPTOR1b*.

These results suggest that the inability to degrade *RAPTOR1b* mRNA via RIDD underlies the hypersensitivity of *ire1ab* mutants to salt stress and further support a functional link between IRE1 and TOR.

## Discussion

Balancing growth and stress responses is essential for organismal survival^22,33,35^. The ER stress sensor IRE1 plays a central role in maintaining this balance^24,54,55^, yet the mechanisms linking IRE1 to growth regulation have remained unclear. Here, we uncover a direct role for IRE1 in suppressing TOR activity through RIDD-mediated degradation of *RAPTOR1b* mRNA. This establishes *RAPTOR1b* as the first RIDD target with a defined function under both physiological and abiotic stress conditions.

TOR promotes growth in response to nutrients^56,57^, thus increasing the burden on ER protein folding^58^. Our findings suggest that IRE1 restricts TOR activity to optimize root growth and prevent proteotoxic stress. We further show that this IRE1-TOR axis intersects with ABA signaling. Specifically, PYL1 inactivation in Col-0 phenocopies the *ire1ab* short-root phenotype, but has no additive effect in *ire1ab*, placing ABA signaling downstream of IRE1 function. Since ABA and auxin jointly regulate root development and auxin has been implicated in ER stress responses^59–65^, future work is needed to determine if disrupted ABA signaling in *ire1ab* also impairs auxin-mediated processes.

Our data demonstrate that IRE1-dependent TOR regulation is independent of the canonical UPR-TFs. The absence of TOR hyperactivation in *bzip28/60*, combined with the hypersensitivity to IRE1 endonuclease inhibition and lethality of the *ire1ab;bzip28* triple mutant^50,66^, supports that RIDD and bZIP28-dependent responses serve compensatory roles in growth and stress tolerance^16–18^.

While RAPTOR1b is known to be regulated post-translationally^67–70^, recent findings have demonstrated that its transcript levels also change dynamically during abiotic stress^29^. Our identification of *RAPTOR1b* as a RIDD target links IRE1 directly to these stress-responsive transcriptional changes, raising the possibility that RIDD contributes to the downregulation of TOR signaling during prolonged stress.

Based on our findings, we propose a model in which IRE1 fine-tunes TOR signaling through RIDD of *RAPTOR1b* mRNA (Figure 6G). Under favorable conditions, this maintains optimal TOR activity and allows growth while limiting ER stress. Upon abiotic stress, ABA activates SnRK2 kinases that phosphorylate RAPTOR1b to further suppress TOR^33^, shifting the balance toward stress adaptation (Figure 6G, left). Without functional IRE1, RAPTOR1b accumulates, TOR becomes hyperactive, and ABA signaling is impaired compromising stress responses and root growth (Figure 6G, right).

In conclusion, we establish RIDD as a critical mechanism by which IRE1 regulates TOR activity and ABA responsiveness, extending the role of IRE1 beyond canonical UPR signaling. These findings provide a foundation for exploring broader roles of RIDD in growth–stress balance and identifying additional targets of IRE1 during environmental challenges.

## Methods

### Plant growth and phenotyping

All plants used in this study were grown under standard long day growth conditions. For most experiments, seedlings were sown on ½ LS plates (1.2% agar) supplemented with 1% sucrose. For experiments using AZD-8055, sucrose was excluded from the media to avoid sucrose-induced stimulation of TOR. For germination experiments, seeds were sown on plates with corresponding treatments and then stratified for 2 days. Once transferred to the growth chamber, photos of plates were taken every 24 hours. Germination rates were calculated my counting the number of seedlings that showed the emergence of the radicle. For root growth, seedlings were grown on vertical plates and photos were taken at day 5, 7, and 10. Root lengths were measured manually with Fiji^71^. T-DNA Knockout mutant lines used in this study include: *ire1ab* (*ire1a,* WISCDSLOX420D09; *ire1b, SAIL_238_F07*), *bzip28/60* (*bzip28,* SALK_132285*; bzip60,* SALK_050203), and *raptor1b* (*raptor1b-1,* SALK_078159).

### Mesh transfer experiments and drug treatment

Square pieces of 100μm Nylon mesh (ELKO Filtering Company) were autoclaved and placed on the surface of solidified ½ LS plates. Seeds were surface sterilized and sown onto the nylon mesh. Seeds were stratified for 2 days and then transferred to a growth chamber. For root growth experiments with ABA, seedlings were allowed to germinate for 3 days and then the mesh was then transferred to fresh plates supplemented with the appropriate drug or hormone allowing for the simultaneous and rapid transfer of many seedlings without the risk of imposing mechanical damage. Experiments using the ABA inhibitor AA1 (Kerafast, Cat. ECR004) were performed with 5μM ABA concentrations to clearly show the effects of the inhibitor. All other experiments with exogenous ABA were performed with 1 or 10μM ABA.

### ActinomycinD treatment

For experiments to check *RAPTOR1b* transcript degradation, Col-0 and *ire1ab* seedlings were sown on large (24x24cm) plates covered with a sterile piece of nylon mesh in three rows and allowed to grow vertically for 10 days under standard growth conditions. On day 10, the plates were flooded with liquid ½ LS supplemented with 25μM ActD or the corresponding volume of DMSO for mock treatment. Plates were returned to the growth chamber and were incubated horizontally for 12 hours. The excess liquid media was removed, and root tips were harvested for RNA extraction. The experiment was repeated for three biological replicates.

### ABA treatment for gene expression

Col-0 and *ire1ab* seedlings were sown on ½ LS plates with 1% sucrose covered with sterile nylon mesh and grown for 10 days. Mesh was transferred to large plates filled with liquid LS supplemented with 10μM ABA or the equivalent volume of ethanol as a mock treatment. Seedlings were returned to the growth chamber and incubated for 6 hours. Liquid media was removed from the plate and 5mm root tips were harvested for RNA extraction.

### gRIDD analysis of Arabidopsis transcriptome

Putative RIDD targets in Arabidopsis were identified using gRIDD to analyze the transcriptome of Arabidopsis seedlings (TAIR10_cdna_20101214_updated) from The Arabidopsis Information Resource (TAIR), Carnegie Institution, Stanford, CA and National Center for Genome Resources, Santa Fe, NM (https://www.arabidopsis.org/download/list?dir=Genes%2FTAIR10_genome_release%2FTAIR10_blastsets).

### RNA isolation and RT-qPCR analysis

Root tips were flash frozen with liquid nitrogen and ground to a fine powder using glass beads and a Retch MM301 (Retch; Haan, Germany) tissue grinder with 4 rounds of shaking at 30hz for 30 seconds each with refreezing between rounds. RNA was purified with the Macherey-Nagel NucleoSpin RNA Plant Kit (Macherey-Nagel) following manufacturers’ instructions. cDNA was generated with the iScript cDNA synthesis kit (BioRad). Gene expression was measured using SYBR detection (Abclonal) and the Applied Biosystems 7500 fast realtime PCR system.

Ubiquitin was used as a housekeeping control. Primers used for this experiment are listed in (Supplemental Table 1). Data points represent averages of three technical replicates. Each experiment was repeated across three independent biological replicates.

### Cloning

Primers were designed to amplify the coding sequence of AtPYL1 with the addition of attB sites for gateway cloning (AT5G46790, Invitrogen Gateway System). The CDS of PYL1 was amplified from cDNA from Col-0 seedlings by PCR. The PCR product was cloned into pDONR207. In order to generate PYL1S199A and S119D, primers were designed to perform site directed mutagenesis PCR. After amplification of the full plasmid, the reactions were DpnI treated to remove all template DNA and then transformed into chemically competent Top10 *E.coli* cells. The sequences of the resulting entry clones were confirmed by sequencing.

Sequence verified clones were then cloned into the pEarlygate104 destination vector by traditional Gateway cloning, sequence verified and transformed into chemically competent Agrobacterium cells (GV3101).

### Protein extraction and S6K phosphorylation assays

For S6K phosphorylation assays, root tip-enriched samples were collected by collecting ∼5mm portions of the root. 60-100 root tips per genotype were collected for each of 3 or 4 technical replicates. The tissue was flash frozen in liquid nitrogen and ground to a fine powder by shaking 5 times at 30Hz/sec for 30 seconds with re-freezing between each run. 50-100ul of protein extraction buffer (PBS, protease inhibitor cocktail (Roche, cOmplete Mini, EDTA-free54), and PhosSTOP (Roche)) was added to the frozen tissue and immediately vortexed until the tissue was thawed. Samples were then transferred to fresh microcentrifuge tubes without glass beads and spun at 21,000rpm for 5, 10, and finally 20 minutes at 4°C. The supernatant was transferred to a fresh tube between runs. Protein concentrations were measured using the BioRad Protein Assay. 5μg of protein was loaded onto 2 12%SDS-PAGE gels. Protein was transferred to nitrocellulose membranes and blocked for a minimum of 1 hour in 3% BSA. Membranes were incubated in primary antibodies against S6K1/2 (Agrisera, AS121855) and pS6K (Abcam, AB207399) overnight at 4° in 3% BSA. Membranes were washed three times for 20 minutes in chilled TBST and incubated in HRP-conjugated Goat anti-rabbit secondary antibody in 2% Nonfat milk in TBST for a minimum of 2 hours. Blots were developed with the SuperSignal™ West Femto Maximum Sensitivity system. Band intensity was calculated with FIJI and ratios of pS6K/S6K1/2 were used to quantify TOR activity.

### Sample preparation and protein extractions for targeted Mass spectrometry

Col-0 and *ire1ab* seeds were sown on the surface of sterilized nylon mesh placed on ½ LS plates supplemented with 1% sucrose. Seeds were stratified for 48 hours and then allowed to germinate for 5 days. In order to have enough total protein for targeted mass spectrometry, root tips from ∼12,000 seedlings per genotype were harvested for each genotype for three independent biological replicates. Each biological replicate consisted of ∼12,000 seedlings plated in two rows across 90 plates. The root tips from all 12,000 seedlings were pooled for protein extractions and flash frozen in liquid nitrogen. Across all three replicates, root tips from approximately 72,000 seedlings were collected for analysis. Crude lysates were prepared as described previously^72^. Briefly, frozen tissue was ground in liquid nitrogen by mortar and pestle and then further ground into a finer powder with the Retch mill. 1mL of protein extraction buffer (4% (wt/vol) SDC and 100 mM Tris-HCl (pH 8.5)) was added to the ground tissue and the samples were incubated at 95°C for 5 minutes. Samples were transferred to fresh tubes without glass beads and were homogenized with sonication (3x 30seconds 1s on, 1s off). 50 μl of each homogenized sample were aliquoted into fresh tubes for protein quantification and the remaining lysates were stored at -80°C until submission. Protein concentrations were quantified using the Pierce™ BCA Protein Assay Kit.

### Recombinant IRE1B_­­_production and in-vitro fluorescence-based RNase assay

The linker and cytosolic region of AtIRE1B was subcloned into the pET-28b(+) vector using the HindIII and XhoI restriction sites and was subsequently transformed into E. coli BL21(DE3) cells as described previously^73^. Single colonies were inoculated in 5 ml of LB containing kanamycin (100 μg/ml) and grown overnight at 37°C. Then 1 ml of overnight culture was inoculated into 50 ml of LB containing kanamycin and grown overnight at 37°C. The following day, 20 ml of culture was then inoculated into 500 ml of LB containing kanamycin and grown to an OD600 of 1.0 at 37°C before being induced with 1mM IPTG. Induced cultures were then cultured overnight at 16°C. Cells were pelleted and stored at -80°C. Cell pellets were lysed by sonication in TALON native binding buffer (50 mM Sodium phosphate pH 7.4, 300 mM NaCl, 1x Roche cOmplete Mini EDTA-free protease inhibitor cocktail) then affinity purified using TALON Metal Affinity Resin (Takara) according to the manufacturers protocol at 4°C, using 10 mM imidazole in wash buffer and eluting with buffer containing 150 mM imidazole. Recombinant IRE1B was buffer exchanged and concentrated into protein storage buffer (50 mM HEPES pH 7.5, 300 mM NaCl, 10% glycerol) using an Amicon Ultra-15 30K to a concentration greater than 35 μM, then aliquoted and stored at -80°C until used for RNase assays. An RNA hairpin probe consisting of a known stem-loop substrate of the human XBP1 RNA, fluorophore, and quencher designed previously (5’ Cy5-CAUGUCCGCAGCGCAUG-BHQ2 3’^47^) was synthesized by IDT for use in the in vitro RNase assays. Assays were performed by assembling two master mixes in assay buffer (25mM HEPES pH 7.5, 5mM MgCl, 150mM NaCl, 5% glycerol), the first containing the hairpin probe and the second containing recombinant IRE1B and additional additives relevant to specific experiments (i.e. ATP/ADP, 4μ8c). The final concentration of the relevant components in the final 150 μl reaction volume were IRE1B: 250 nM; ADP 250nM; ATP 250nM; RNA probe 320 nM; 4μ8c 0-2000 nM. All reactions were initiated simultaneously by combining the two master mixes using a multichannel pipette in a 96 well plate. RNA cleavage was measured kinetically by increasing Cy5 fluorescence intensity over time.

### IRE1B-mRNA modeling

Predicted dimeric structure of IRE1B (UniProt: F4KH40_ARATH) together with the RNA sequences of interest were generated via AlphaFold 3.0.1^49^. UniProt lists three separate protein IDs for IRE1B (Q93VJ2_ARATH, F4KH40_ARATH, and F4KH41_ARATH). We selected the sequence associated with F4KH40 for our models because this is the entry that is directly linked to TAIR. Q93VJ2_ARATH and F4KH40_ARATH are very similar sequences with F4KH40_ARATH possessing six additional amino acids in the N-terminal portion of the protein.

The presence of these six amino acids explains why our model highlights N826 rather than the previously reported N820 as the critical amino acid for IRE1B nuclease activity^50^.

F4KH41_ARATH is an uncharacterized potential splice variant of IRE1B that is 20 amino acids shorter than F4KH40_ARATH. Rather than using the default of 5 conformations from the same seed, we sampled more of the available conformations using 3 different seeds and 5 conformations generated from each seed for a total of 15 IRE1B-RNA complexes for each RNA sequence of interest. Typical ranking scores for the models were about 0.4 on a scale of 0-1, reflective of the large unstructured regions usually present in the predicted structures (Fig. 6A). The protein-RNA complexes were visualized in VMD 1.9.4a58^74^. Purpose-built python scripts leveraging the VMD python API as well as matplotlib^75^, scipy^76^, and numpy^77^ analyzed specific structural statistics from the predicted structures to quantify observations from visual inspection of the complexes.

### In vitro RIDD assay

His-tagged IRE1B_­­_was expressed in E. coli BL21 cells, induced with 0.3 mM IPTG at OD600 0.6-0.8, and grown at 18°C for 16 hours. Harvested cells were lysed by ultrasonication in lysis buffer (50 mM phosphate, pH 8.0, 300 mM NaCl, 2 mM PMSF). The clarified supernatant was incubated with Ni-NTA Agarose Resin (GoldBio), washed sequentially with lysis buffer containing 10 mM to 60 mM imidazole, and eluted with lysis buffer containing 250 mM imidazole. Pooled fractions were buffer-exchanged into storage buffer (HEPES pH 7.5, 300 mM NaCl, 10% glycerol) using Amicon® Ultra Centrifugal Filters (30 kDa MWCO). Isolated protein was stored at -20°C.

5μg of total RNA was incubated in assay buffer (25mM HEPES pH 7.5, 5mM MgCl, 150mM NaCl, 5% glycerol) with and without 250nM IRE1B_­­_and 250nM ATP for three hours at 22°C. Additional reactions with heat-inactivated IRE1B_­­_were used as a negative control.1μg of RNA from each assay was used for cDNA synthesis with the iScript cDNA synthesis kit (BioRad).

Transcript levels of RAPTOR1b were measured by qRT-PCR using SYBR detection (Abclonal) and the Applied Biosystems 7500 fast realtime PCR system. Ubiquitin was used as a housekeeping control. Unspliced bZIP60 was used as a positive control for IRE1B_­­_activity and LST8-1 was used as a negative control for RIDD.

### Mass spectrometry

Mass spectrometry to detect the phosphorylated form of AtPYL1 was performed by the Proteomics Core facility at MSU as described below:

#### Proteolytic Digestion

Protein samples were mixed with 100mM Tris-HCl (pH 8.5) supplemented to 4% (w/v) sodium deoxycholate (SDC) and then reduced and alkylated by adding TCEP and chloroacetamide at 10mM and 40mM, respectively. Samples were incubated for 5min at 45C with shaking at 2000 rpm in an Eppendorf ThermoMixer C. Trypsin, in 50mM ammonium bicarbonate, was added at a 1:100 ratio (wt/wt) and the mixture was incubated at 37C overnight with shaking at 1500 rpm in the Thermomixer. Final volume of each digest was ∼300uL.

#### Phosphopeptide Isolation

Digests were subjected to phosphopeptide isolation according to the method of Humphrey, et.al. Isolated phosphopeptides were then desalted using SDB-RPS (3M Empore) StageTips, dried by vacuum centrifugation and stored at -20C.

#### LC/MS/MS Analysis

Isolated phosphopeptides were re-suspended in 2%ACN/0.1%TFA to 13uL. Injections of 10uL were automatically made using a Thermo (www.thermo.com) EASYnLC 1200 onto a Thermo Acclaim PepMap RSLC 0.1mm x 20mm C18 trapping column and washed for ∼5min with buffer A. Bound peptides were then eluted over 125min onto a Thermo Acclaim PepMap RSLC 0.075mm x 500mm resolving column with a gradient of 5%B to 25%B at 90min, ramping to 42%B at 114min, to 100% B at 115min and held at 100%B for the duration of the run (Buffer A = 99.9% Water/0.1% Formic Acid, Buffer B = 80% Acetonitrile/0.1% Formic Acid/19.9% Water) at a constant flow rate of 300nl/min. Column temperature was maintained at a constant temperature of 50°C using and integrated column oven (PRSO-V2, Sonation GmbH, Biberach, Germany). Eluted peptides were sprayed into a ThermoScientific Q-Exactive HF-X mass spectrometer (www.thermo.com) using a FlexSpray spray ion source and peptides of interest were assayed using a Parallel Reaction Monitoring method.

#### Data Analysis

The resulting MS/MS spectra are converted to peak lists Mascot Distiller, v2.8.3 (www.matrixscience.com) and searched against a protein sequence database containing target protein sequences appended with common laboratory contaminants (downloaded from www.thegpm.org, cRAP project) or against all protein entries available in the TAIR, v10, reference proteome (downloaded from www.arabidopsis.org) using the Mascot2 searching algorithm, v 2.8.0.1. The Mascot output was then analyzed using Scaffold, v5.2.2 (www.proteomesoftware.com) to probabilistically validate protein identifications. Assignments validated using the Scaffold 1%FDR confidence filter are considered true. Targeted spectra were interpreted using Skyline.

*Mascot parameters for all databases were as follows:*

- Allow up to 2 missed tryptic sites
- Fixed modification of Carbamidomethyl Cysteine,
- Variable modification of oxidation of Methionine, Phosphorylation of Serine, Threonine
- Peptide tolerance of +/- 10ppm
- MS/MS tolerance of 0.02 Da
- FDR calculated using randomized database search

## Acknowledgements

This study was supported primarily by the National Institutes of Health (R35GM136637) to FB with contributing support from National Institutes of Health (R35GM155317) to JVV, Chemical Sciences, Geosciences and Biosciences Division, Office of Basic Energy Sciences, Office of Science, US Department of Energy (award number DE-FG02-91ER20021) to FB and JVV, and MSU AgBioResearch (MICL02598) to FB. This work was also supported in part through computational resources and services provided by the Institute for Cyber-Enabled Research at Michigan State University. JVV receives partial salary support from MSU AgBioResearch. We also thank the Research Technology Support Facility Proteomics Core at Michigan State University for assistance with the targeted mass spectrometry experiments and data analysis.

## Author Contributions

B.C.R and F.B. conceptualized the project and designed the experiments; B.C.R., E.R.A, and J.Y.K. performed the experiments; J.V.V. generated and models and interpreted modeling results. B.C.R. and F.B. interpreted the results; B.C.R. and F.B. wrote the manuscript.

---

**Supplemental Figure 1: TOR-dependent phosphorylation site in PYL1 is highly phosphorylated in *ire1ab* and the site is conserved across the PYL family.** a. The sequence of the target peptide standard used to detect phosphorylated PYL1 and a representative spectrum demonstrating complete coverage of the phospho-peptide by mass spectrometry. Colors represent the b- and y-ions identified in the fragmentation data. b. Quantification of the Total TIC values for the phosphopeptide in Col-0 and ire1ab root tips across three biological replicates. c. Multiple sequence alignment demonstrating that the phosphorylation site (red box) is conserved across all 14 members of the Arabidopsis PYL family. The phosphopeptide sequence used for targeted mass spectrometry is highlighted in blue.

Supplemental Figure 2: 4μ8C is a potent inhibitor of the nuclease activity of Arabidopsis IRE1B. **a-c.** *In vitro* RNAse assays with purified IRE1B_­­_demonstrating the effects of increased substrate concentration (a) and ADP/ATP (b) on the RNase activity of IRE1B *in vitro*. IRE1B’s RNAse activity is inhibited by 4μ8C in a concentration-dependent manner (c). d. Representative images of Col-0 and *ire1ab* seedlings grown on plates supplemented with increasing concentrations of 4μ8C exhibit reduced primary root growth. Red dashed lines represent the average root length of each genotype under each treatment. **e**. Average relative root length (4μ8C /Mock) of Col-0 and *ire1ab* seedlings grown at different concentrations of 4μ8C. Dashed line represents the average length of untreated *ire1ab* roots.

**Supplemental Figure 3: AlphaFold3 predicts that IRE1B recognizes the putative consensus sequence in *RAPTOR1b* mRNA. a.** Example AlphaFold3 output for the IRE1B dimer complexed with *RAPTOR1b* mRNA. The colors in this representation indicate AlphaFold3 confidence scores, with bluer colors indicating lower confidence and redder colors indicating higher confidence (see the associated scale bar). The IRE1B dimer is largely confidently predicted, particularly the KEN domain shown at the “top” of the protein where the mRNA interacts. **b**. Difference in the θ and η angles along the mRNA backbone. The gray region indicates the residues with different sequences between the WT and mutant, with the most noticeable structural changes occurring just upstream of this region. **c.** Difference in SASA between the nucleobase component of the mRNA sequence between WT and mutant. The gray region again indicates the residues with different sequences between the WT and mutant. **d.** Representative snapshot highlighting which nucleobases (spheres) are often flipped out from the overall mRNA structure (blue cartoon) to make contact with the protein environment, such as N826 in the KEN domain (green cartoon). The highlighted residue and nucleobases are colored according to atom identity, with nitrogens colored in blue, oxygens in red, and carbons either yellow (protein) or gray (nucleobase) depending on their identity. Generally, the nucleobases that are flipped out are further along in the sequence for the WT compared with the mutant.

**Supplemental Figure 4: The RIDD consensus sequence in AtRAPTOR1b is conserved across multiple plant species.** Multiple sequence alignment of RAPTOR1B sequences demonstrating that the consensus sequence is conserved across plant species. The Arabidopsis sequence is highlighted in orange. Exact matches for the sequence are highlighted in yellow. Other colors represent sequences where the consensus sequence contains a base substitution at the second (gray), fifth (blue), or second and fifth (peach) position within the consensus sequence. These positions correspond to the previously reported consensus variants, Var2 and Var5^47^.

**Supplemental** Figure 5: *ire1ab* is hypersensitive to salt stress. **a.** Representative images of seedlings grown on plates supplemented with 50mM or 100mM NaCl. Red dashed lines represent the average root length of each genotype under each treatment. **b.** Quantification of relative primary root length (NaCl/Mock) demonstrates that *ire1ab* seedlings are hypersensitive to salt stress.

**Supplemental Table 1: Primers used for cloning and qRT-PCR in this study.**

**Supplemental Data 1: List of potential RIDD targets identified via gRIDD software. Supplemental Data 2: Direct File for AlphaFold3 models of IRE1B and RNA**

