## Supplemental Figures for "IRE1 Regulates TOR Signaling via RIDD of *RAPTOR1b* to Coordinate Growth and Stress Adaptation"

### Supplemental Figure and Supplementary information

1. **Supplemental Figure 1:** TOR-dependent phosphorylation site in PYL1 is highly phosphorylated in *ire1ab* and the site is conserved across the PYL family.
2. **Supplemental Figure 2:** 4μ8C is a potent inhibitor of the nuclease activity of Arabidopsis IRE1B.
3. **Supplemental Figure 3:** AlphaFold3 predicts that IRE1B recognizes the putative consensus sequence in *RAPTOR1b* mRNA.
4. **Extended Data 4:** The RIDD consensus sequence in *AtRAPTOR1b* is conserved across multiple plant species.
5. **Supplemental Figure 5**: The *ire1ab* mutant is hypersensitive to salt stress.
6. ***Supplementary Table 1:*** *Primer list*

7. ***Supplementary Data 1:*** *List of genes identified via gRIDD.*
8. ***Supplementary Data 1:*** *Direct File for AlphaFold3 models of IRE1b and RNA.*

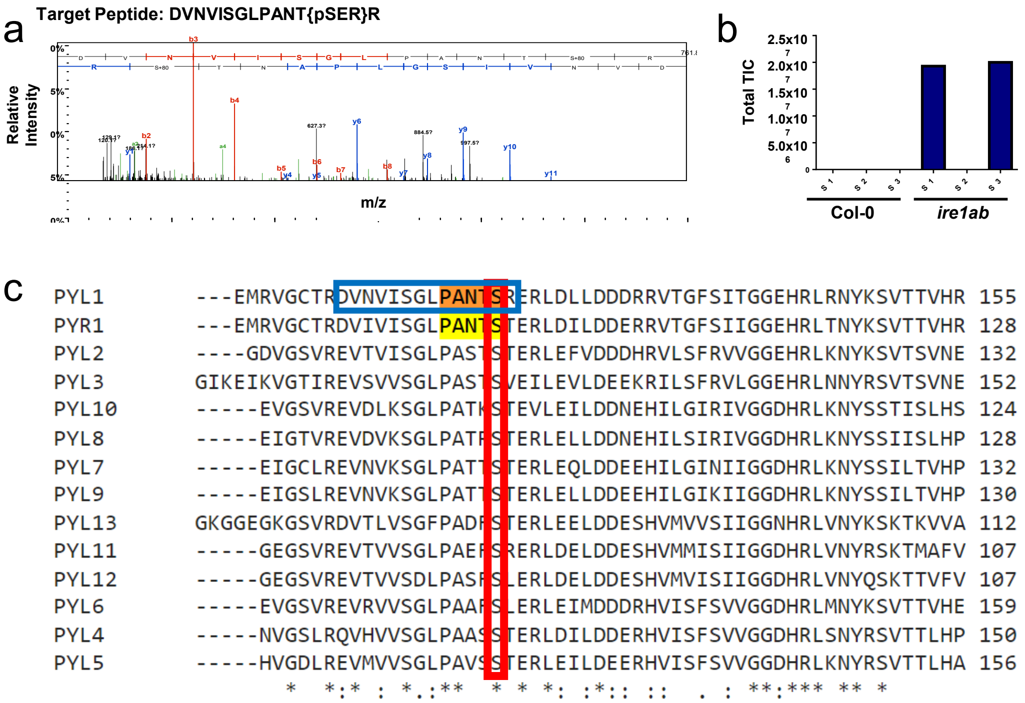


**Supplemental Figure 1: TOR-dependent phosphorylation site in PYL1 is highly phosphorylated in *ire1ab* and the site is conserved across the PYL family. a.** The sequence of the target peptide standard used to detect phosphorylated PYL1 and a representative spectrum demonstrating complete coverage of the phospho-peptide by mass spectrometry. Colors represent the b- and y-ions identified in the fragmentation data. **b.** Quantification of the Total TIC values for the phosphopeptide in Col-0 and ire1ab root tips across three biological replicates. **c.** Multiple sequence alignment demonstrating that the phosphorylation site (red box) is conserved across all 14 members of the Arabidopsis PYL family. The phosphopeptide sequence used for targeted mass spectrometry is highlighted in blue.

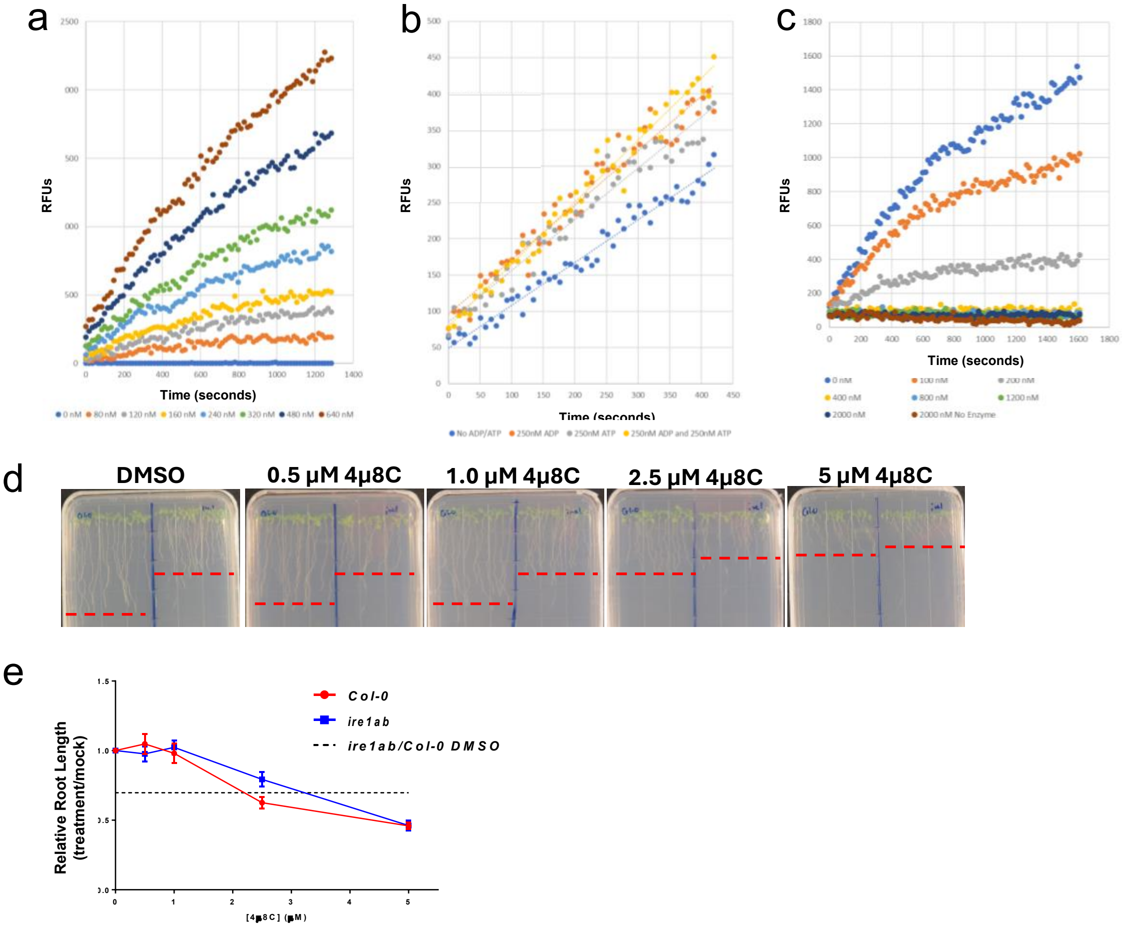


**Supplemental Figure 2:** 4μ8C is a potent inhibitor of the nuclease activity of Arabidopsis IRE1B. **a-c.** *In vitro* RNAse assays with purified IRE1B$_{cyto}$ demonstrating the effects of increased substrate concentration (a) and ADP/ATP (b) on the RNase activity of IRE1B *in vitro*. IRE1B's RNAse activity is inhibited by 4μ8C in a concentration-dependent manner (c). d. Representative images of Col-0 and *ire1ab* seedlings grown on plates supplemented with increasing concentrations of 4μ8C exhibit reduced primary root growth. Red dashed lines represent the average root length of each genotype under each treatment. **e**. Average relative root length (4μ8C /Mock) of Col-0 and *ire1ab* seedlings grown at different concentrations of 4μ8C. Dashed line represents the average length of untreated *ire1ab* roots.

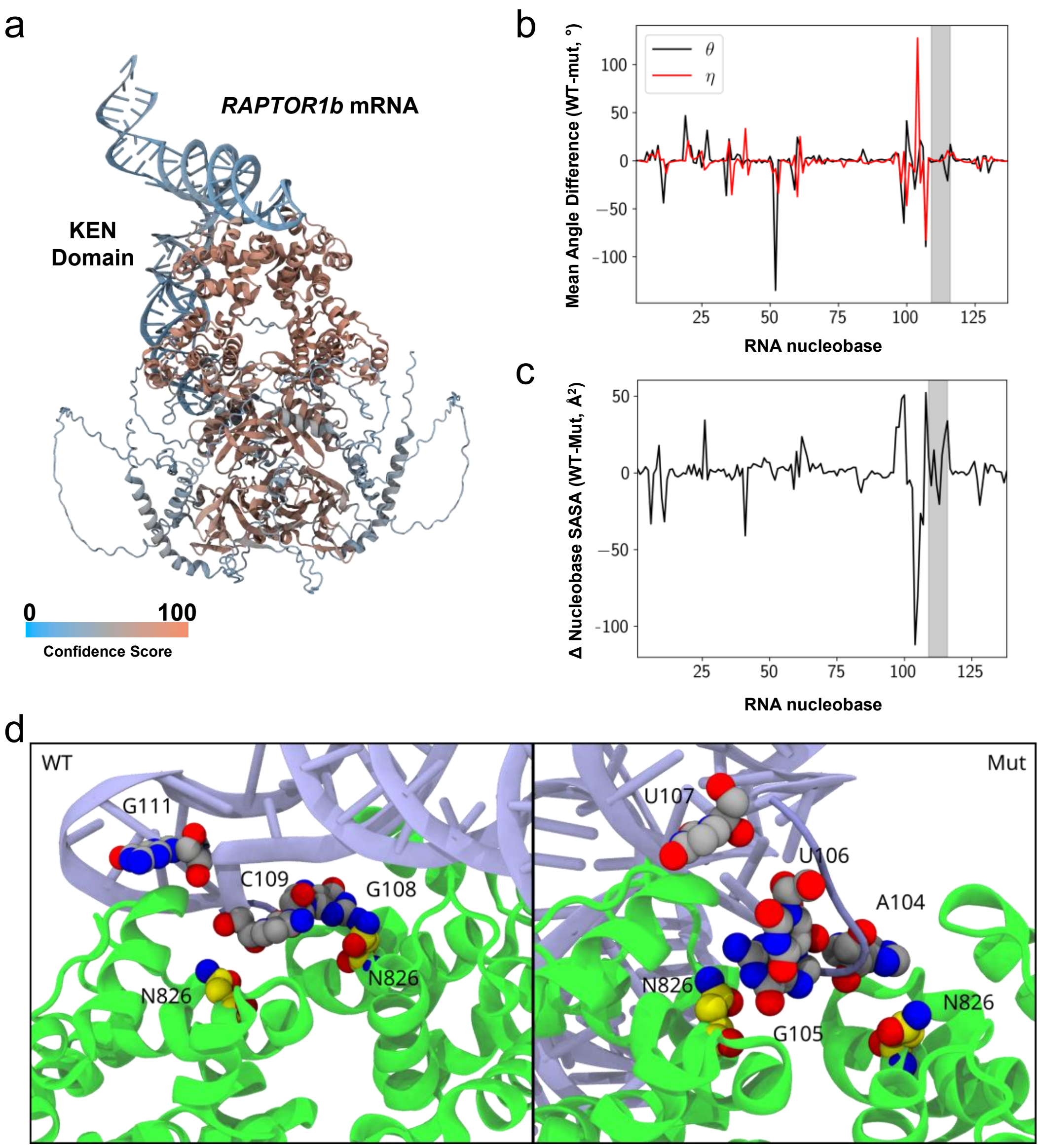


**Supplemental Figure 3: AlphaFold3 predicts that IRE1B recognizes the putative consensus sequence in *RAPTOR1b* mRNA. a.** Example AlphaFold3 output for the IRE1B dimer complexed with *RAPTOR1b* mRNA. The colors in this representation indicate AlphaFold3 confidence scores, with bluer colors indicating lower confidence and redder colors indicating higher confidence (see the associated scale bar). The IRE1B dimer is largely confidently predicted, particularly the KEN domain shown at the "top" of the protein where the mRNA interacts. **b**. Difference in the θ and η angles along the mRNA backbone. The gray region indicates the residues with different sequences between the WT and mutant, with the most noticeable structural changes occurring just upstream of this region. **c.** Difference in SASA between the nucleobase component of the mRNA sequence between WT and mutant. The gray region again indicates the residues with different sequences between the WT and mutant. **d.** Representative snapshot highlighting which nucleobases (spheres) are often flipped out from the overall mRNA structure (blue cartoon) to make contact with the protein environment, such as N826 in the KEN domain (green cartoon). The highlighted residue and nucleobases are colored according to atom identity, with nitrogens colored in blue, oxygens in red, and carbons either yellow (protein) or gray (nucleobase) depending on their identity. Generally, the nucleobases that are flipped out are further along in the sequence for the WT compared with the mutant.

Identified Consensus sequence: CNGCAGN

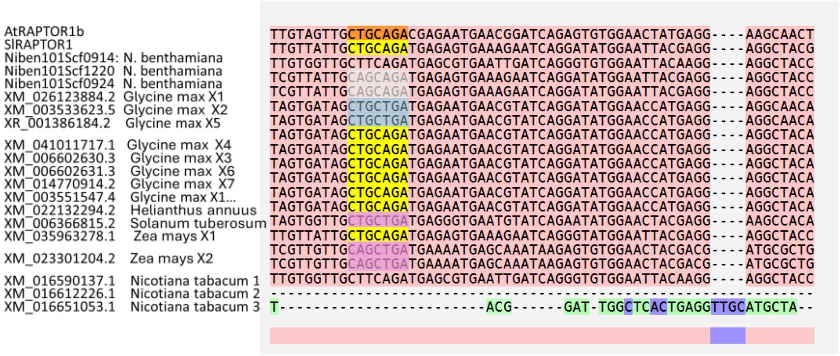


**Supplemental Figure 4: The RIDD consensus sequence in AtRAPTOR1b is conserved across multiple plant species.** Multiple sequence alignment of RAPTOR1B sequences demonstrating that the consensus sequence is conserved across plant species. The Arabidopsis sequence is highlighted in orange. Exact matches for the sequence are highlighted in yellow. Other colors represent sequences where the consensus sequence contains a base substitution at the second (gray), fifth (blue), or second and fifth (peach) position within the consensus sequence. These positions correspond to the previously reported consensus variants, Var2 and Var5[47].

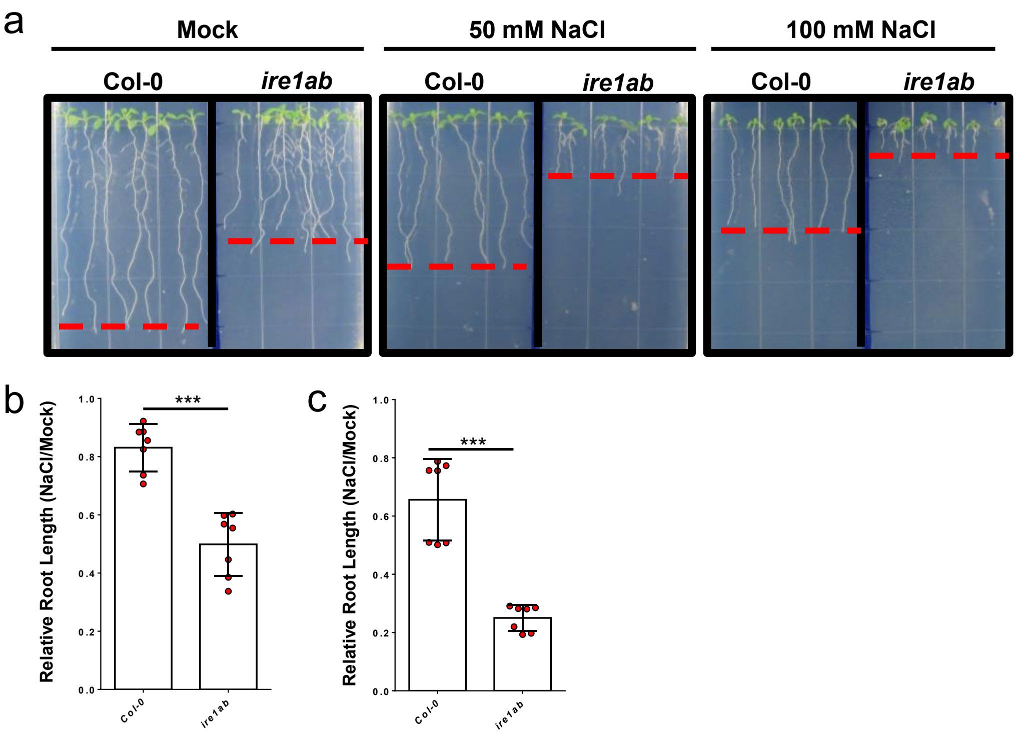


**Supplemental Figure 5:** *ire1ab* is hypersensitive to salt stress. **a.** Representative images of seedlings grown on plates supplemented with 50mM or 100mM NaCl. Red dashed lines represent the average root length of each genotype under each treatment. **b.** Quantification of relative primary root length (NaCl/Mock) demonstrates that *ire1ab* seedlings are hypersensitive to salt stress.
