## Supplemental Data 2 for "IRE1 Regulates TOR Signaling via RIDD of *RAPTOR1b* to Coordinate Growth and Stress Adaptation": countcontacts.py

```
#The idea behind this script is to count in how many frames the mutated sequence is at least somewhat near GLN 826.
import glob
from vmd import Molecule, atomsel
import numpy as np
def loadcifs(d):
	ciflist = glob.glob(d + "/*/*cif")
	print(ciflist)
	mol = Molecule.Molecule()
	for cif in ciflist:
		mol.load(cif, filetype="pdbx")
	return mol
def evaluatecontacts(mol):
	psel = atomsel("protein and resid 826") #This is the ASN implicated in catalysis.
	nsel = atomsel("sequence CGGCGGAU or sequence CUGCAGAC") #This is the mutant or WT sequence
	contactexists = np.empty(mol.numFrames(), dtype=bool)
	for f in range(mol.numFrames()):
		psel.frame = f
		nsel.frame = f
		if len(psel.contacts(nsel, 5)[0]):
			contactexists[f] = True
		else:
			contactexists[f] = False
		#print(f, psel.contacts(nsel, 10))
	return contactexists
def measurenucleobasesasa(mol):
	nsel = atomsel("nucleic") #This is the mutant or WT sequence
	nside = atomsel("nucleic and not backbone")
	sasa = np.empty((mol.numFrames(), len(list(set(nsel.resid)))))
	for f in range(mol.numFrames()):
		nsel.frame = f
		nside.frame = f
		sasa[f] = nsel.sasaperresidue(1.4, restrict=nside)
		#print(f, psel.contacts(nsel, 10))
	return sasa
l = ['mut', 'wt']
for i, d in enumerate(["../outputs/fullmut", "../outputs/fullwt"]):
	mol = loadcifs(d)
	contactbools = evaluatecontacts(mol)
	sasa = measurenucleobasesasa(mol)
	np.save("sasa-%s.npy" % (l[i]),sasa)
	print(contactbools, np.sum(contactbools))
exit()
```
