## Supplemental Data 2 for "IRE1 Regulates TOR Signaling via RIDD of *RAPTOR1b* to Coordinate Growth and Stress Adaptation": plotpseudotorsions.py

```
import matplotlib.pyplot as plt
plt.rcParams['text.usetex'] = True
plt.rcParams['text.latex.preamble'] = r'\usepackage{sfmath}'
plt.rcParams['font.size'] = 18

# plt.rcParams.update({
#     "text.usetex": True,
#     "font.family": "Helvetica",
#     "text.latex.preamble": [
#     r'\usepackage{tgheros}',    # helvetica font
#     r'\usepackage{sfmath}',   # math-font matching  helvetica
#     r'\sansmath'                # actually tell tex to use it!
#     ]
# })
import numpy as np
from scipy.stats import circmean
fig, ax = plt.subplots(1,1)
l = ['wt', 'mut']
for i in range(len(l)):
	data = np.load("psuedotorsions-%s.npz" % l[i])
	thetas = data['thetas']
	etas = data['etas']
	x = np.array(range(len(thetas)))
	ax.plot(x+1, circmean(thetas,high=180, low=-180,axis=1), label = "%s $\\theta$" % l[i])
	ax.plot(x+2, circmean(etas,high=180, low=-180,axis=1), label = "%s $\\eta$" % l[i])
ax.legend()
ax.set_xlabel("Resid")
ax.set_ylabel("Angle ($^{\\circ}$)")
fig.tight_layout()
fig.savefig("psuedotorsionplot.png", dpi=300)

fig, ax = plt.subplots(1,1)
l = ['wt', 'mut']
wtdata = np.load("psuedotorsions-wt.npz")
wtthetas = wtdata['thetas']
wtetas = wtdata['etas']
mutdata = np.load("psuedotorsions-mut.npz")
mutthetas = mutdata['thetas']
mutetas = mutdata['etas']
x = np.array(range(len(wtthetas)))
ax.plot(x+1, circmean(wtthetas-mutthetas,high=180, low=-180,axis=1), label = "$\\theta$")
ax.plot(x+2, circmean(wtetas-mutetas,high=180, low=-180,axis=1), label = "$\\eta$")
ax.legend()
ax.set_xlabel("Resid")
ax.set_ylabel("Angle Difference ($^{\\circ}$)")
fig.tight_layout()
fig.savefig("psuedotorsionplotchange.png", dpi=300)

fig, ax = plt.subplots(1,1)
l = ['wt', 'mut']
wtdata = np.load("psuedotorsions-wt.npz")
wtthetas = wtdata['thetas']
wtetas = wtdata['etas']
mutdata = np.load("psuedotorsions-mut.npz")
mutthetas = mutdata['thetas']
mutetas = mutdata['etas']
x = np.array(range(len(wtthetas)))
ax.plot(x+1, (((circmean(wtthetas,high=180, low=-180,axis=1)-circmean(mutthetas,high=180, low=-180,axis=1))+180)%360)-180, label = "$\\theta$", color='k')
ax.plot(x+2, (((circmean(wtetas,high=180, low=-180,axis=1)-circmean(mutetas,high=180, low=-180,axis=1))+180)%360)-180, label = "$\\eta$", color='r')
ax.legend()
ax.set_xlim(left=1,right=np.amax(x)+2)
ax.axvspan(109,116, alpha=0.2,color="k")
ax.set_xlabel("RNA Residue")
ax.set_ylabel("Mean Angle Difference (WT-mut, $^{\\circ}$)")
fig.tight_layout()
fig.savefig("psuedotorsionplotchange2.png", dpi=300)
exit()
```
