## Supplemental Data 2 for "IRE1 Regulates TOR Signaling via RIDD of *RAPTOR1b* to Coordinate Growth and Stress Adaptation": plotsasa.py

```
import matplotlib.pyplot as plt
plt.rcParams['text.usetex'] = True
plt.rcParams['text.latex.preamble'] = r'\usepackage{sfmath}'
plt.rcParams['font.size'] = 18

# plt.rcParams.update({
#     "text.usetex": True,
#     "font.family": "Helvetica",
#     "text.latex.preamble": [
#     r'\usepackage{tgheros}',    # helvetica font
#     r'\usepackage{sfmath}',   # math-font matching  helvetica
#     r'\sansmath'                # actually tell tex to use it!
#     ]
# })
import numpy as np
from scipy.stats import circmean
fig, ax = plt.subplots(1,1)
l = ['wt', 'mut']
for i in range(len(l)):
	data = np.load("sasa-%s.npy" % l[i])
	
	x = np.array(range(len(data[0])))
	ax.plot(x+1, np.mean(data,axis=0),label=l[i].upper())
ax.axvspan(109,116, alpha=0.2,color="k")
ax.legend()
ax.set_xlim(left=1,right=np.amax(x)+1)
ax.set_xlabel("RNA Residue")
ax.set_ylabel("Mean Nucleobase SASA (Å$^2$)")
fig.tight_layout()
fig.savefig("sasaplot.png", dpi=300)

fig, ax = plt.subplots(1,1)
wtdata = np.load("sasa-wt.npy")
mutdata = np.load("sasa-mut.npy")
x = np.array(range(len(data[0])))
ax.plot(x+1, np.mean(wtdata,axis=0)-np.mean(mutdata,axis=0),color='k')
ax.axvspan(109,116, alpha=0.2,color="k")
ax.set_xlim(left=1,right=np.amax(x)+1)
ax.set_xlabel("RNA Residue")
ax.set_ylabel("$\\Delta$ Nucleobase SASA (WT-Mut, Å$^2$)")
fig.tight_layout()
fig.savefig("sasaplotdiff.png", dpi=300)
```
