## Supplemental Data 2 for "IRE1 Regulates TOR Signaling via RIDD of *RAPTOR1b* to Coordinate Growth and Stress Adaptation": pseudotorsions.py

```
#The idea behind this script is to count in how many frames the mutated sequence is at least somewhat near GLN 826.
import glob
from vmd import Molecule, atomsel, measure
import numpy as np
def loadcifs(d):
	ciflist = glob.glob(d + "/*/*cif")
	print(ciflist)
	mol = Molecule.Molecule()
	for cif in ciflist:
		mol.load(cif, filetype="pdbx")
	return mol
#https://nakb.org/basics/nucleotides.html#VIRTUAL/
def pseudotorsion(mol):
	#nsel = atomsel("name P C4' and (sequence CGGCGGAU or sequence CUGCAGAC)") #This is the mutant or WT sequence
	nsel = atomsel("name P C4'") #Sequence is from resid 109 to 116
	print(len(nsel), nsel.name, nsel.resid)
	idxs = nsel.index
	thetas = np.zeros(((len(idxs)-4)//2,mol.numFrames()))
	etas = np.zeros(((len(idxs)-4)//2,mol.numFrames()))
	for i in range(0, len(idxs[:-4]), 2):
		#First theta is for resid 1
		thetas[i//2] = measure.dihedral(idxs[i+0], idxs[i+1], idxs[i+2], idxs[i+3], first=0, last=15)
		#First eta is for resid 2
		etas[i//2] = measure.dihedral(idxs[i+1], idxs[i+2], idxs[i+3], idxs[i+4], first=0, last=15)
	return (thetas, etas)
l = ['mut', 'wt']
for i, d in enumerate(["../outputs/fullmut", "../outputs/fullwt"]):
	mol = loadcifs(d)
	thetas, etas = pseudotorsion(mol)
	np.savez("psuedotorsions-%s.npz" % (l[i]), thetas=thetas, etas=etas)
exit()
```
