## Supplemental Data 2 for "IRE1 Regulates TOR Signaling via RIDD of *RAPTOR1b* to Coordinate Growth and Stress Adaptation": runalphafold3.sh

```
#!/bin/bash
#SBATCH --time=4:00:00
#SBATCH --gres=gpu:1
#SBATCH -C [neh|nel|nal|nif|nvf] ## We want the good GPUs
#SBATCH --cpus-per-task=8
#SBATCH --mem=48G
#SBATCH -o outputs/fullmutinference.out


nvidia-smi
#For inference, we need sufficiently new GPUs, or at least a workaround.
#See https://github.com/google-deepmind/alphafold3/issues/59 for working with the V100s
COMPUTECAP=`nvidia-smi --query-gpu=compute_cap --format=csv | tail -n1`

EXPECTCAP='8.0'
printf -v versions '%s
%s' "$COMPUTECAP" "$EXPECTCAP"
if [[ $versions = "$(sort -V <<< "$versions")" ]]; then
    echo 'FAILURE'
    export XLA_FLAGS="--xla_disable_hlo_passes=custom-kernel-fusion-rewriter"
    export xlaargs="--flash_attention_implementation=xla"
else
    echo 'SUCCESS'
    export xlaargs=""
fi

cd $SLURM_SUBMIT_DIR
singularity exec      --nv      --bind $SLURM_SUBMIT_DIR/inputs:/root/af_input      --bind $SLURM_SUBMIT_DIR/outputs:/root/af_output      --bind /mnt/gs21/scratch/vermaasj/Alphafold3:/root/models      --bind /mnt/gs21/scratch/vermaasj/alphafolddatabases:/root/public_databases      /mnt/gs21/scratch/vermaasj/Alphafold3/alphafold3.sif      python /root/models/run_alphafold.py      --norun_data_pipeline      --json_path=/root/af_output/fullmut/fullmut_data.json      --model_dir=/root/models      --db_dir=/root/public_databases      --output_dir=/root/af_output      --force_output_dir $xlaargs
```
