## Supplemental Data 2 for "IRE1 Regulates TOR Signaling via RIDD of *RAPTOR1b* to Coordinate Growth and Stress Adaptation": runicer.py

```
import glob
import numpy as np
import os
import time
import subprocess
inputlist = sorted(glob.glob("inputs/full*json"))

#This is if you want to do everything in one shot. This is a bad idea, since you are tying up a GPU while doing the multiple sequence alignment.
script = '''#!/bin/bash
#SBATCH --time=04:00:00
#SBATCH --gres=gpu:1
#SBATCH -C [neh|nel|nal|nif|nvf|nvl|nfh] ## We want the good GPUs
#SBATCH --cpus-per-task=8
#SBATCH --mem=48G
#SBATCH -o outputs/%s.out

cd $SLURM_SUBMIT_DIR
singularity exec \
     --nv \
     --bind $SLURM_SUBMIT_DIR/inputs:/root/af_input \
     --bind $SLURM_SUBMIT_DIR/outputs:/root/af_output \
     --bind /mnt/gs21/scratch/vermaasj/Alphafold3:/root/models \
     --bind /mnt/gs21/scratch/vermaasj/alphafolddatabases:/root/public_databases \
     /mnt/gs21/scratch/vermaasj/Alphafold3/alphafold3.sif \
     python /root/models/run_alphafold.py \
     --json_path=/root/af_input/%s \
     --model_dir=/root/models \
     --db_dir=/root/public_databases \
     --output_dir=/root/af_output 

'''
#This is the typical first step. *most* of the time it'll finish in 4 hours, but sometimes it won't.
sequencealignonly = '''#!/bin/bash
#SBATCH --time=12:00:00
#SBATCH --cpus-per-task=8
#SBATCH --mem=48G
#SBATCH -o outputs/%s.out

cd $SLURM_SUBMIT_DIR
singularity exec \
     --nv \
     --bind $SLURM_SUBMIT_DIR/inputs:/root/af_input \
     --bind $SLURM_SUBMIT_DIR/outputs:/root/af_output \
     --bind /mnt/gs21/scratch/vermaasj/Alphafold3:/root/models \
     --bind /mnt/gs21/scratch/vermaasj/alphafolddatabases:/root/public_databases \
     /mnt/gs21/scratch/vermaasj/Alphafold3/alphafold3.sif \
     python /root/models/run_alphafold.py \
     --norun_inference \
     --json_path=/root/af_input/%s \
     --model_dir=/root/models \
     --db_dir=/root/public_databases \
     --output_dir=/root/af_output 

'''


#This is the second step. Usually finishes in 4 hours, except for very large proteins.
inferenceonly = '''#!/bin/bash
#SBATCH --time=4:00:00
#SBATCH --gres=gpu:1
#SBATCH -C [neh|nel|nal|nif|nvf] ## We want the good GPUs
#SBATCH --cpus-per-task=8
#SBATCH --mem=48G
#SBATCH -o outputs/%s.out


nvidia-smi
#For inference, we need sufficiently new GPUs, or at least a workaround.
#See https://github.com/google-deepmind/alphafold3/issues/59 for working with the V100s
COMPUTECAP=`nvidia-smi --query-gpu=compute_cap --format=csv | tail -n1`

EXPECTCAP='8.0'
%s
if [[ $versions = "$(sort -V <<< "$versions")" ]]; then
    echo 'FAILURE'
    export XLA_FLAGS="--xla_disable_hlo_passes=custom-kernel-fusion-rewriter"
    export xlaargs="--flash_attention_implementation=xla"
else
    echo 'SUCCESS'
    export xlaargs=""
fi

cd $SLURM_SUBMIT_DIR
singularity exec \
     --nv \
     --bind $SLURM_SUBMIT_DIR/inputs:/root/af_input \
     --bind $SLURM_SUBMIT_DIR/outputs:/root/af_output \
     --bind /mnt/gs21/scratch/vermaasj/Alphafold3:/root/models \
     --bind /mnt/gs21/scratch/vermaasj/alphafolddatabases:/root/public_databases \
     /mnt/gs21/scratch/vermaasj/Alphafold3/alphafold3.sif \
     python /root/models/run_alphafold.py \
     --norun_data_pipeline \
     --json_path=/root/af_output/%s/%s_data.json \
     --model_dir=/root/models \
     --db_dir=/root/public_databases \
     --output_dir=/root/af_output \
     --force_output_dir $xlaargs

'''
for element in inputlist:
	name = element[7:-5]
	print(name)
	rerun = False
	if os.path.exists("outputs/%s/%s_model.cif" % (name.lower(), name.lower())):
		print(name + " exists!")
	else:
		rerun = True
	if os.path.exists("outputs/%s/ranking_scores.csv" % (name.lower())):
		data = np.loadtxt("outputs/%s/ranking_scores.csv" % (name.lower()), delimiter=",", skiprows=1,usecols=2)
		print(data)
		if np.mean(data) < 0:
			rerun = True
	rerun=True
	if rerun:
		fout = open("runalphafold3.sh", "w")
		if os.path.exists("outputs/%s/%s_data.json" % (name.lower(), name.lower())):
			fout.write(inferenceonly % (name + "inference", "printf -v versions '%s\n%s' \"$COMPUTECAP\" \"$EXPECTCAP\"",name.lower(), name.lower()))
		else:
			fout.write(sequencealignonly % (name, element[7:]))
		fout.close()
		#exit()
		subprocess.call("sbatch --job-name %s runalphafold3.sh" % name, shell=True)
		#time.sleep(2) #Wait for 2 seconds between submissions. I'm really depending on the batch system making a bunch of temporary files upon submission.
```
