## Supplemental Data 2 for "IRE1 Regulates TOR Signaling via RIDD of *RAPTOR1b* to Coordinate Growth and Stress Adaptation": runinputs.sh

```
#!/bin/bash
PWD=`pwd`
inputs=`ls inputs`
echo $inputs
for input in $inputs
do
     echo $input
singularity exec \
     --nv \
     --bind $PWD/inputs:/root/af_input \
     --bind $PWD/outputs:/root/af_output \
     --bind /tank/alphafold3:/root/models \
     --bind /tank/alphafolddatabases:/root/public_databases \
     /tank/alphafold3.sif \
     python /root/models/run_alphafold.py \
     --json_path=/root/af_input/$input \
     --model_dir=/root/models \
     --db_dir=/root/public_databases \
     --output_dir=/root/af_output \
     --num_diffusion_samples=20 \
     --num_seeds=3

done
```
