## Supplementary figures and images for "IRE1 Regulates TOR Signaling via RIDD of *RAPTOR1b* to Coordinate Growth and Stress Adaptation"

### psuedotorsionplot.png

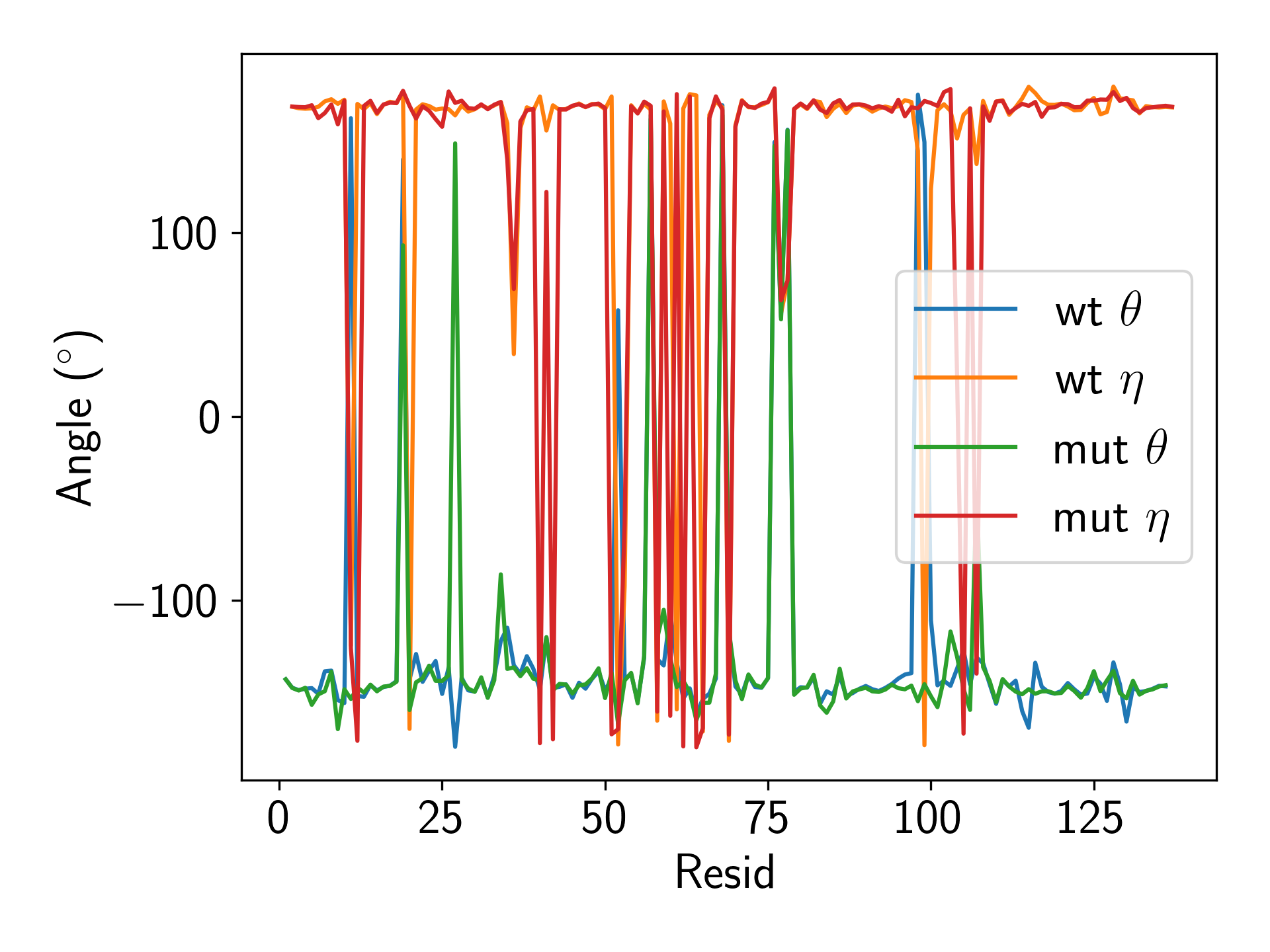

### psuedotorsionplotchange2.png

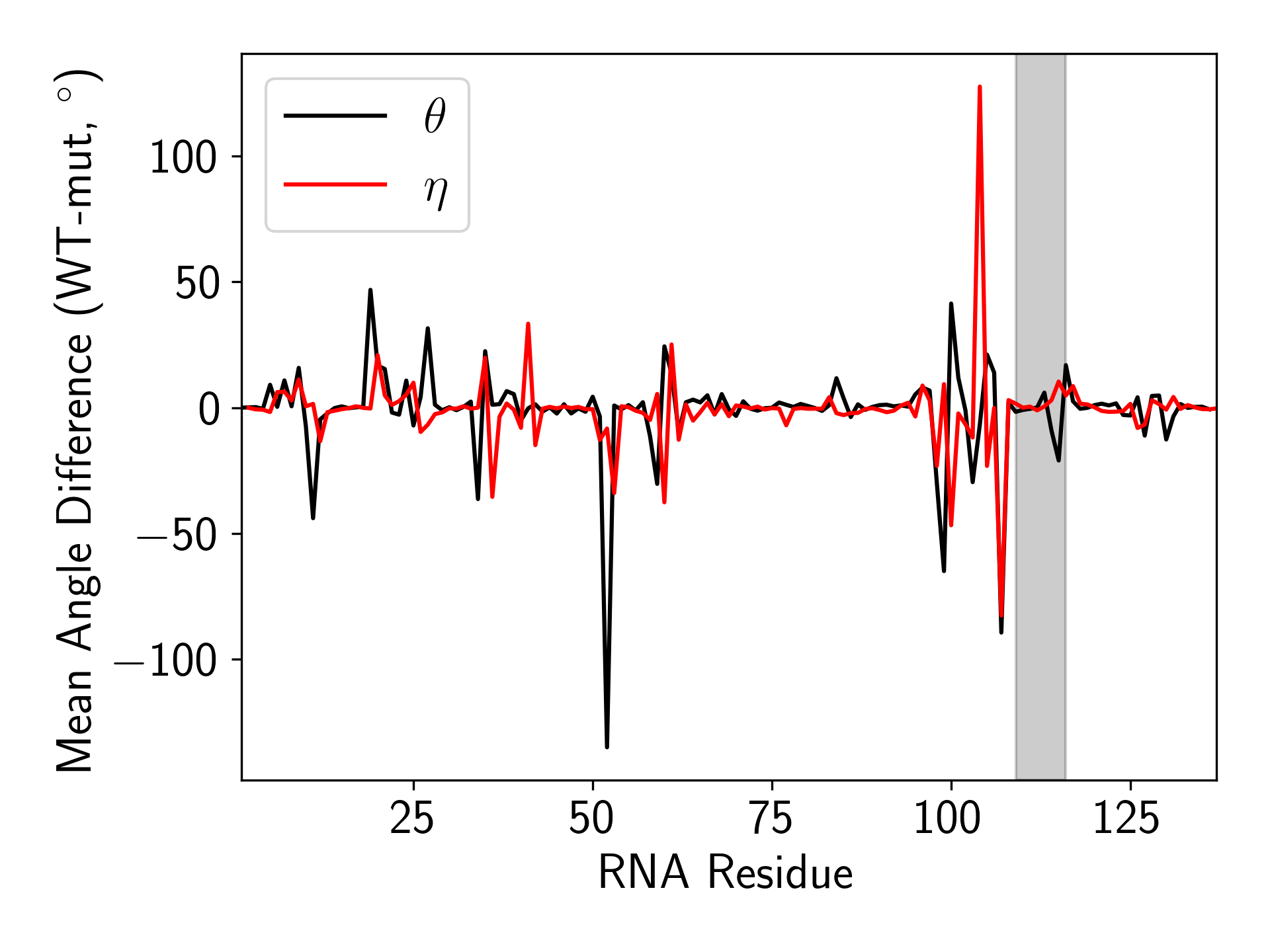

### psuedotorsionplotchange.png

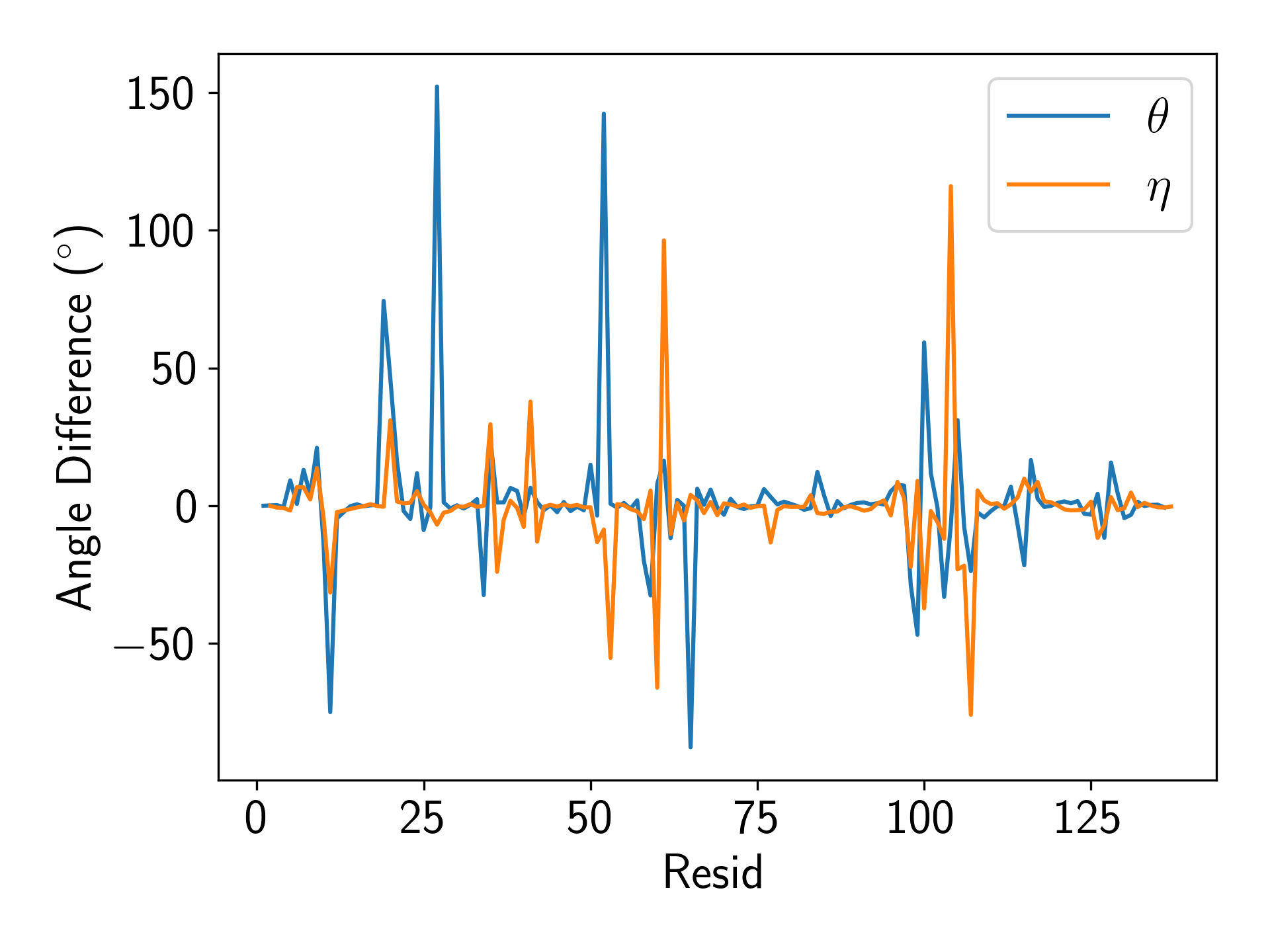

### sasaplot.png

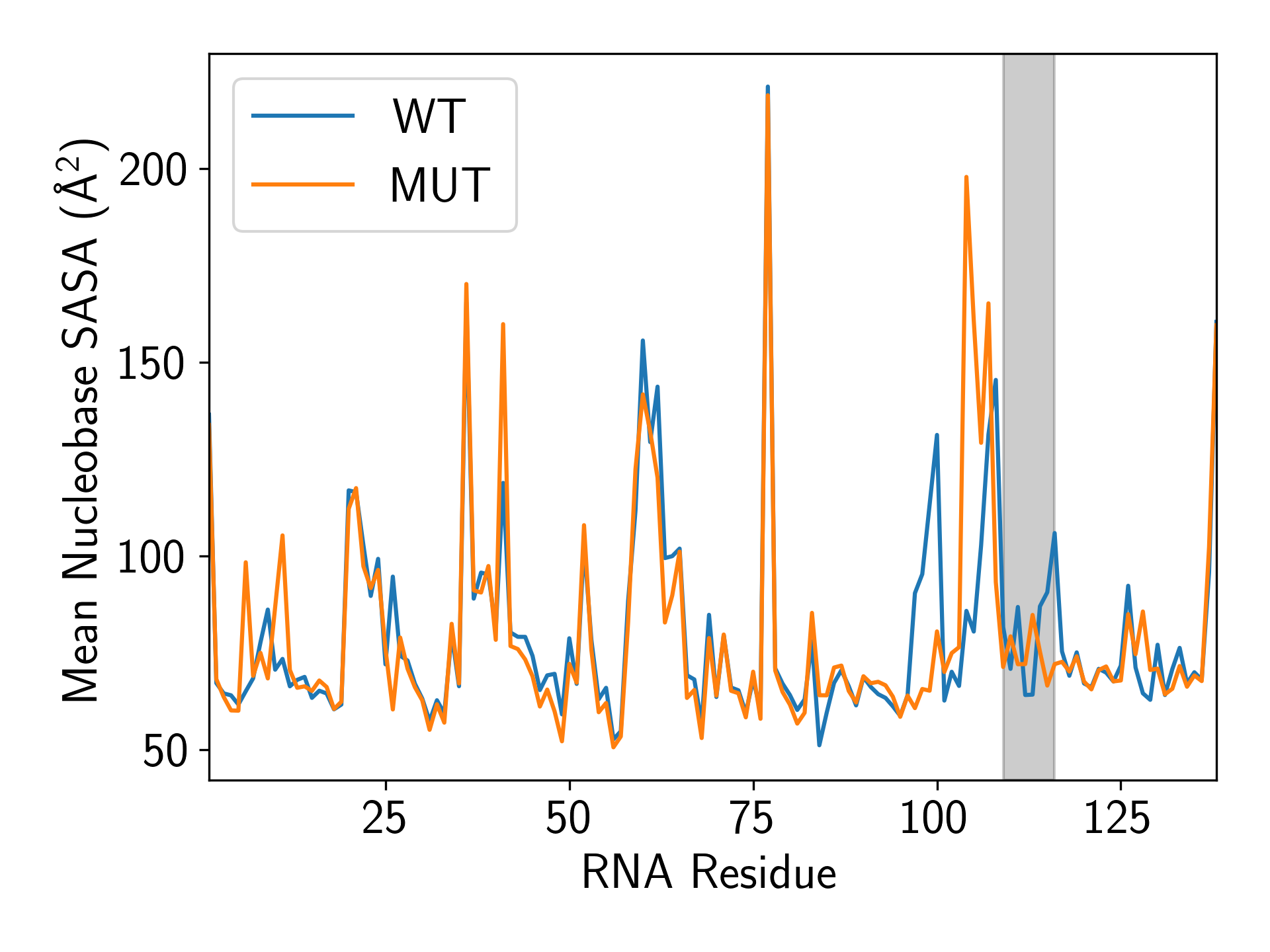

### sasaplotdiff.png

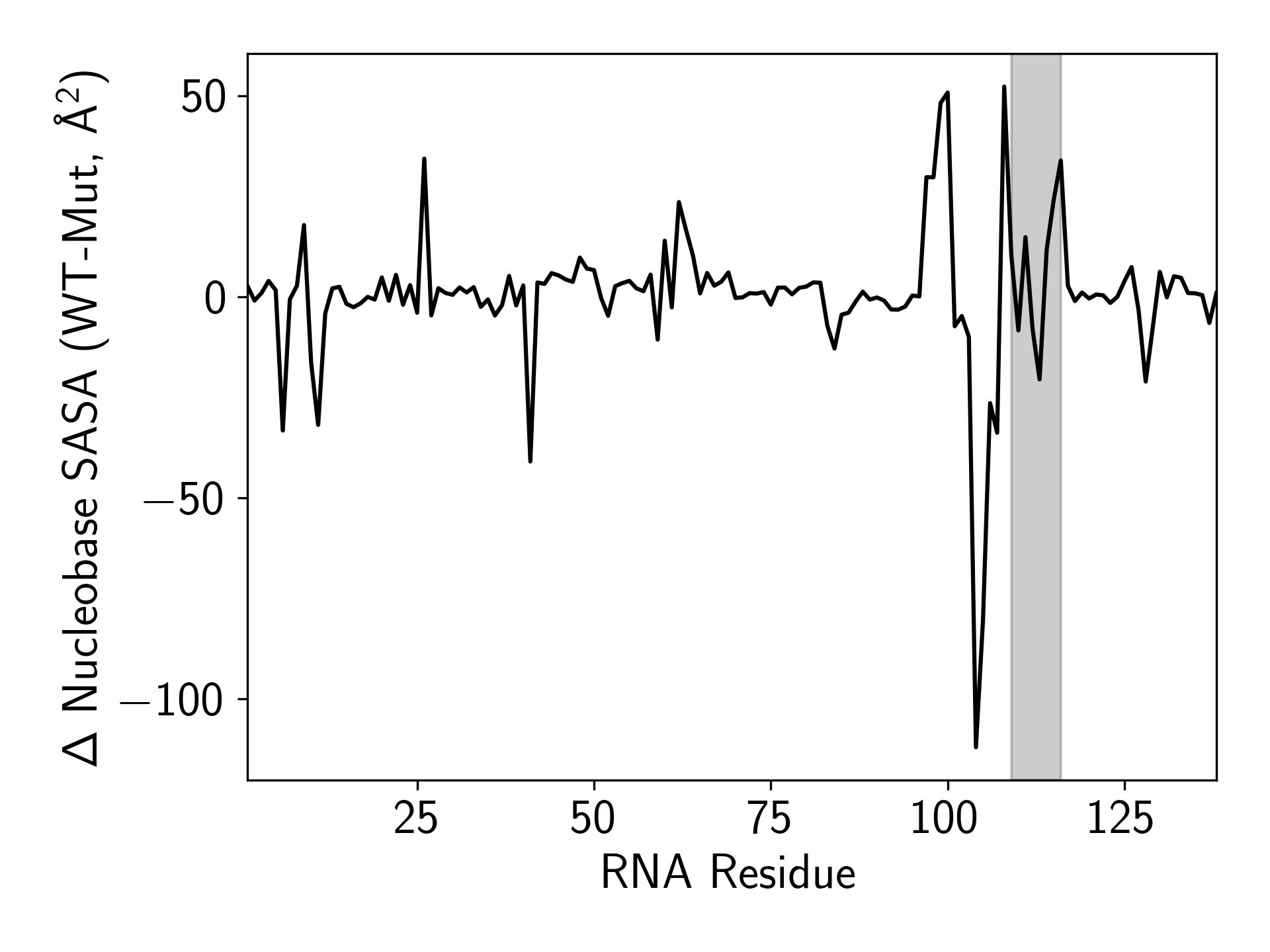
