## Supplemental Table 1 for "IRE1 Regulates TOR Signaling via RIDD of *RAPTOR1b* to Coordinate Growth and Stress Adaptation"

| Target Gene | **Primer Type** | Primer Sequence |
| --- | --- | --- |
| Ubiquitin | **qRT-PCR Forward** | GGCCTTGTATAATCCCTGATGAATAAG |
| Ubiquitin | **qRT-PCR Reverse** | AAAGAGATAACAGGAACGGAAACATAGT |
| RD29A | **qRT-PCR Forward** | GGAAGTGAAAGGAGGAGGAGGAA |
| RD29A | **qRT-PCR Reverse** | CACCACCAAACCAGCCAGATG |
| RD29B | **qRT-PCR Forward** | GAATCAAAAGCTGGGATGGA |
| RD29B | **qRT-PCR Reverse** | TGCTCTGTGTAGGTGCTTGG |
| RAB18 | **qRT-PCR Forward** | GGCTTGGGAGGAATGCTTCA |
| RAB18 | **qRT-PCR Reverse** | CGCTTGAGCTTGACCAGACT |
| RAPTOR1b | **qRT-PCR Forward** | GCCAACTGGGATACAAGGTTTG |
| RAPTOR1b | **qRT-PCR Reverse** | AGTTCCACACTCTGATCCGTTC |
| LST8-1 | **qRT-PCR Forward** | GGATGGAGAATTTCTTGTAACAGC |
| LST8-1 | **qRT-PCR Reverse** | TGATGACCTTGGTACACTTTCAC |
| PYL1 | **Cloning Forward + attB site** | ACAAGTTTGTACAAAAAAGCAGGCTCCATGGCGAATTCAGAGTCCTCCTCCT |
| PYL1 | **Cloning Reverse + attB site** | ACCACTTTGTACAAGAAAGCTGGGTTTACCTAACCTGAGAAGAGTTGTTGTT |
| PYL1S119A | **Site directed mutagenesis forward** | AATACGGCGCGAGAGAGATTAGATCTGT |
| PYL1S119A | **Site directed mutagenesis reverse** | TCTCTCGCGCCGTATTCGCCGGTAATCC |
| PYL1S119D | **Site directed mutagenesis forward** | CGGCGAATACGGATCGAGAGAGATTAGATCT |
| PYL1S119D | **Site directed mutagenesis reverse** | TCTCTCGATCCGTATTCGCCGGTAATCCACT |
| pDONR207 | **Sequencing** | TCGCGTTAACGCTAGCATGGATCTC |
| pDONR207 | Sequencing | GTAACATCAGAGATTTTGAGACAC |

**Supplementary Table 1: Primer list**
